# Subcellular distribution of PP1 isoforms in holoenzyme complexes

**DOI:** 10.1101/2022.09.09.507380

**Authors:** Virja Mehta, Delphine Chamousset, Jennifer Law, Sarah Ooi, Denise Campuzano, Vincent Nguyen, François-Michel Boisvert, Greg B. Moorhead, Laura Trinkle-Mulcahy

## Abstract

Unlike its counterpart Ser/Thr kinases, the predominant Ser/Thr protein phosphatase 1 (PP1) is a promiscuous enzyme that gains its subcellular localization and substrate specificity from a large panel of regulatory proteins with which it associates in predominantly dimeric complexes. Inhibition of specific PP1-mediated dephosphorylation events relies on targeting the regulatory rather than the catalytic subunit, which in turn relies on a comprehensive understanding of the holoenzyme complexes that underlie its distribution throughout the cell. Proteomic, bioinformatic and biochemical screens have assembled lists of putative regulatory proteins, which have been studied to varying degrees. We took a non-biased approach to link steady-state localization to complexes, using a combination of fluorescence imaging, cellular fractionation and quantitative affinity purification/mass spectrometry (AP/MS) to map interactomes for PP1ɑ/β/ɣ in 3 human cell lines. Comparing the distribution of each isoform between the pool of identified regulatory subunits highlighted key signaling pathways and identified c20orf27 as a novel PP1 regulatory protein. Steady-state association of a large fraction of PP1 with the evolutionarily conserved SDS22 was demonstrated, as was redistribution at the entry to mitosis. This is consistent with recent work suggesting that SDS22 acts as a PP1 sink from which it can be recruited as needed. Moving forward, this approach can be used to assess the redistribution of PP1 during other cellular processes or in response to perturbations or disease states, facilitating identification of the relevant complexes and the design of strategies to target them therapeutically.

## Introduction

Reversible protein phosphorylation, the most common post-translational modification (PTM), involves the covalent attachment of phosphate groups to amino acid residues by protein kinases and their subsequent removal by protein phosphatases. This acts as a molecular switch that can modulate protein conformation and/or protein-protein interactions, leading to alterations in enzymatic activity, subcellular localization, turnover of targets and signaling by other PTMs. Phosphoregulation plays a role in most cellular processes, including signaling, migration, cell cycle progression and metabolism, and is a key therapeutic target in diseases in which these processes are deregulated. Recent advances in mass spectrometry (MS)-based phosphoproteomics revealed that the majority of cellular proteins can be phosphorylated, and most on multiple sites[1]. Whereas thousands of these phosphorylation sites have been linked to specific upstream kinases, only ∼7% have been mapped to a cognate phosphatase[2]. Mapping each phosphatase to its physiological substrates is an ongoing challenge, as it is essential for understanding the specific roles that they play in complex phospho-signaling networks. The predominant phosphorylated amino acid is Serine (Ser), which accounts for >80% of phospho-events[3]. Threonine (Thr) and Tyrosine (Tyr) account for the bulk of the remaining phosphosites, with phosphoryation also demonstrated to a lesser extent on other amino acid residues[4]. Although the human genome encodes a similar number of Tyr phosphatases to counteract Tyr kinase activity, Ser/Thr kinases outnumber Ser/Thr phosphatases by a factor of 10. This difference reflects a broader *in vitro* substrate specificity for the phosphatases, with *in vivo* specificity provided by additional regulatory mechanisms that govern their localization and activity. The two most abundant Ser/Thr phosphatases are classified as either type 1 (PP1) or type 2 (PP2). They were originally defined using biochemical assays and then further divided based on their requirements for metal ions and substrate specificity[5]. Both families have been linked to the regulation of signaling pathways throughout the cell, with PP1 in particular playing a prominent role in nuclear events[5],[6]. Most Ser/Thr phosphatases, including PP1 and PP2A, are regulated and achieve their substrate specificity through the association of the catalytic subunit with a range of regulatory or “targeting” subunits[7]. This results in the combinatorial generation of a large and diverse group of multimeric PP1 and PP2A holoenzyme complexes, each with its own subset of substrates and mechanism(s) of regulation. The distribution of these complexes thus determines the subcellular distribution of catalytic activity at any given time.

The PP1 catalytic subunit (PP1_cat_), which is ubiquitously expressed in all eukaryotic cells, is estimated to account for 40-70% of Ser/Thr dephosphorylation events[8]. Mammalian PP1_cat_ is found primarily as three isoforms PP1 (ɑ, β/δ, γ) that are encoded by three distinct genes, with additional tissue-specific splicing of PP1γ generating the testis-enriched and sperm-specific isoform PP1γ2[9]. The PP1c isoforms are >89% identical in amino acid sequence, with minor variations primarily at their NH2 and COOH termini[10]. Loss of function and biochemical studies of individual PP1 genes in eukaryotic organisms suggest some level of compensation but also highlight distinct phenotypes associated with the disruption of a single gene[9].

Despite their abundance, PP1 catalytic subunits are not found as free monomers in eukaryotic cells[11]. This is not surprising, as a free subunit would be capable of dephosphorylating a large population of phosphoproteins[12]. Instead, there is a molar excess of the PP1 regulatory proteins that compete for binding to a limited pool of endogenous PP1[7]. The three PP1_cat_ isoforms show distinct and dynamic localization patterns in cells that reflect underlying differences in their distribution between regulatory complexes[13]–[15]. Overexpression of a single regulatory subunit can override these distinct localization patterns[15], highlighting the importance of validating functionally relevant interactions and dephosphorylation events under steady-state conditions.

To date, >200 confirmed PP1 interacting proteins have been identified using proteomic and biochemical approaches and yeast two-hybrid and bioinformatic screens[15]–[21]. A broad in silico screen based on a stringent definition of the amino acid “RVxF” motif that binds PP1, for example, included follow-up biochemical approaches to validate association with PP1[22]. The majority of known PP1 regulatory proteins contain an RVxF docking motif, which mediates association with a hydrophobic channel in PP1_cat_[23]. Binding at this site is mutually exclusive (i.e. only one regulatory subunit can bind PP1_cat_ at a time via this motif) and does not influence enzymatic activity, as the catalytic site is 20 Å away[24]. Several PP1 regulatory proteins contain additional PP1-binding sequences, such as SILK and MyPhoNE motifs, that enhance binding and contribute to isoform preference (see [7] for review). There is also a subset of inhibitory PP1 interactors that do not contain a typical RVxF docking motif (e.g. Inh2, SDS22) and can associate with PP1 bound to another regulatory subunit in trimeric complexes[7].

We previously reported the establishment and characterization of HeLa and U2OS cell lines stably expressing PP1 isoforms as fusions with the green fluorescent protein GFP[15],[17],[25]. When expressed stably at low levels, GFP-tagged PP1 isoforms are valid markers for endogenous pools of PP1, permitting direct comparison of their subcellular distributions by live imaging and efficient recovery from cell lysates for interactome mapping[15]. This circumvents both the notorious unreliability of PP1 isoform-specific antibodies for immunostaining and immunoprecipitation[25],[26] and the potential to disrupt the physiological balance of PP1 holoenzyme activity by transient or induced overexpression. Using these cell lines, along with additional HeLa and MCF7 stable cell lines, we set out to map the predominant PP1 holoenzyme complexes that underlie the observed steady-state localization patterns observed for each isoform. To do this, we combined cellular fractionation with quantitative SILAC-based affinity purification/mass spectrometry (AP/MS) and applied stringent criteria to identify high confidence interactors. This revealed both distinct and overlapping roles for the isoforms in signaling complexes, highlighted conserved regulatory and functional characteristics and identified novel complexes. Furthermore, we demonstrated how this approach can be used to detect functionally relevant changes in the subcellular distribution of the catalytic subunit between regulatory complexes, highlighting changes that occur at the transition to M-phase.

## Results

### PP1 isoforms are equally distributed between the cytoplasm and nucleus

Given that PP1 activity exists in a balance in cells, and that the majority of published studies by necessity rely on overexpression of either a catalytic or regulatory subunit to assess their functional interactions, we first assessed the subcellular localization of the endogenous isoforms by proteomic and Western blot analysis. Combining cell fractionation with quantitative MS allowed us to directly compare the subcellular distribution of Ser/Thr protein phosphatases between the cytoplasm (CP) and nucleus (NUC). U2OS cells were differentially labeled with “Light” (L) or “Heavy” (H) SILAC media, followed by fractionation and preparation of CP and NUC extracts. Combining cell equivalent volumes of Heavy CP and Light NUC extract for LC-MS/MS analysis and quantifying H:L ratios for all identified proteins enabled the direct comparison of their distribution between the two compartments (Figure 1a). The >1100 proteins that were identified/quantified (Supplemental Data File 1) included the three PP1 isoforms, PP5, plus several PP2 family members.

**Fig. 1.**
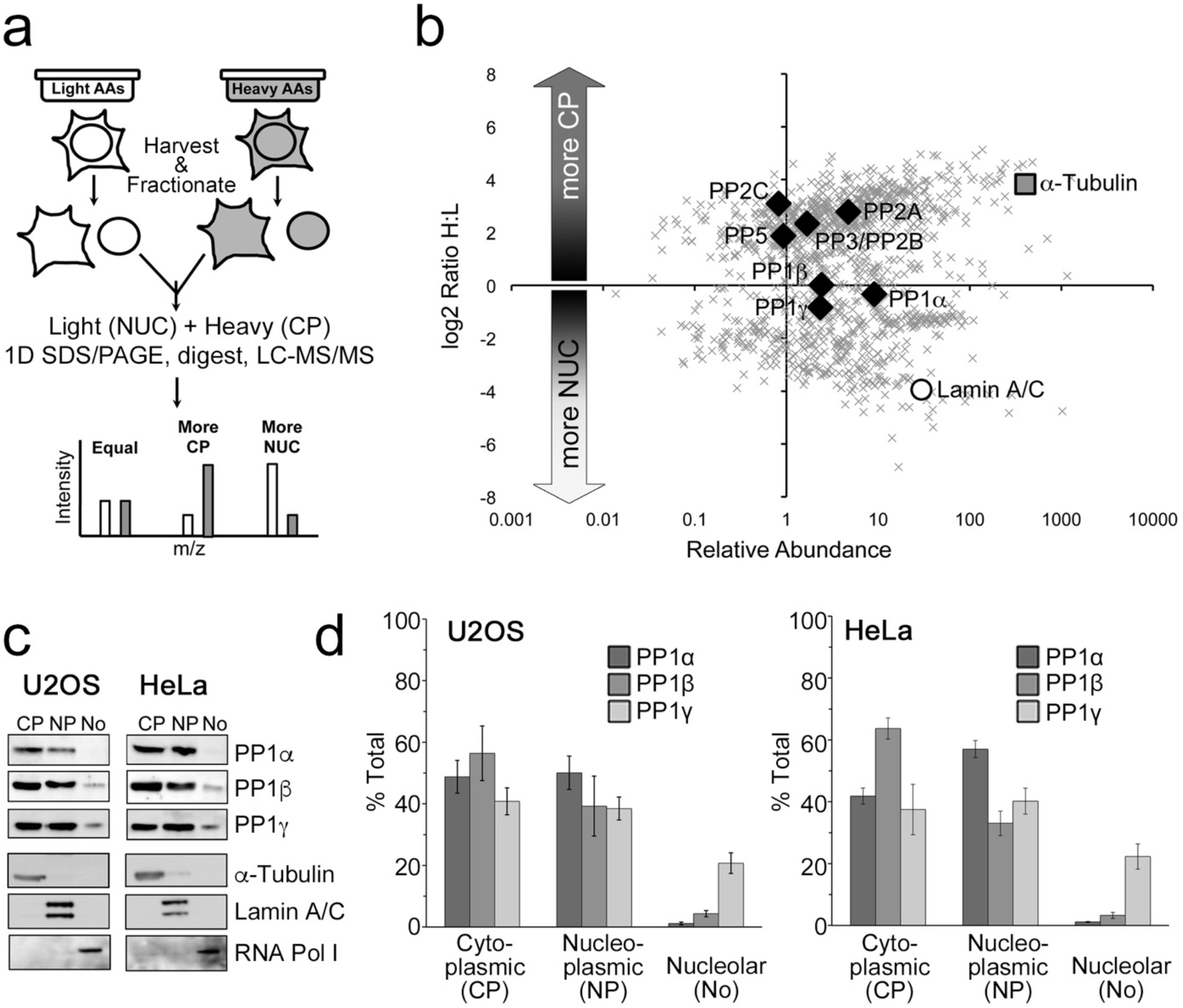
PP1 isoforms exhibit a more equal cytoplasmic/nuclear distribution compared to PP5 and PP2 family serine/threonine phosphatases. (**a**) Design of quantitative subcellular proteome mapping experiment directly comparing the distribution of proteins between cell equivalent volumes of Nuclear (NUC; R0K0 Light media) and Cytoplasmic (CP; R10K8 Heavy media) extracts prepared from U2OS cells. (**b**) Plotting log2 H:L ratio (CP:NUC distribution) vs. relative abundance (summed and normalized peptide intensities) for each identified protein highlights the distribution of the three PP1 isoforms relative to PP5 and PP2 family Ser/Thr phosphatases. Known CP (alpha-tubulin) and NUC (Lamin A/C) markers distribute as expected, confirming the fractionation efficiency. (**c**) Stringently validated PP1 isoform-specific antibodies were utilized for Western blot analysis to assess the distribution of PP1ɑ, β and ɣ between CP, Nucleoplasmic (NP) and Nucleolar (No) extracts from both U2OS and HeLa cells. CP (a-tubulin), NP (Lamin A/C) and No (RNA Pol I; A190) markers confirmed fractionation efficiency. (**d**) Quantitation of the data in **c**, plotted as mean ± SE for three biological replicates.

The full dataset was plotted as log2 Ratio H:L vs. Relative Abundance (Figure 1b), with a log value of 0 representing equal distribution (i.e. H:L Ratio of 1:1) between CP and NUC. We used a threshold of 1 log above/below this value (>2-fold enrichment) to classify proteins as CP-enriched (53%), NUC-enriched (33%) or equally distributed (14%). For reference, the commonly utilized CP marker, α-tubulin, was strongly CP-enriched (log2 Ratio H:L = 3.82), while the commonly utilized NUC marker, Lamin A/C, was strongly NUC-enriched (log2 Ratio H:L = -3.94). Other protein families were found enriched in the expected fractions, such as the CP myosins and filament proteins and the NUC histones and mRNA splicing factors. The distribution of identified Ser/Thr phosphatases between the two subcellular fractions is highlighted on the graph (Figure 1B) and the full list of peptides that were detected for each provided in Supplemental Datal File 1. PP5 and all PP2 family phosphatases that were detected (PP2Aɑ/β, PP2Bɑ, PP2Cɑ/PPM1G) were enriched >2-fold in the CP fraction, as were several PP2A regulatory subunits (PPP2R1A/1B, PPP2R2A, PPP2R4, PPP2R5C/E). In contrast, the PP1 isoforms were found to be more equally distributed, with log2 H:L Ratios of -0.37, 0.014 and -0.84, for PP1α, PP1β and PP1β, respectively.

We next compared the distribution of the endogenous PP1 isoforms between cytoplasmic (CP), nucleoplasmic (NP) and nucleolar (No) fractions in both U2OS and HeLa cells by Western blot analysis, using our in-house PP1 isoform-specific antibodies (Fig. 1c). These show minimal cross-reactivity in Western blot assays, provided the anti-PP1ɑ antibody is first cross-absorbed with the PP1β immunogenic peptide[27]. The intensity per area (minus background) of the isoform band in each subcellular fraction was measured and represented as a fraction of the summed intensity per area of all the fractions (Figure 1D). The distribution was consistent with the CP/NUC patterns observed by MS analysis, and with our previous observations that PP1ɣ is the predominant nucleolar isoform[17].

### GFP-PP1 isoforms stably expressed at low levels are active phosphatases that form holoenzyme complexes and show similar subcellular distributions to the endogenous proteins

Our previous PP1 interactome mapping had focused on the α and γ isoforms, as we found that the β isoform was more susceptible to degradation. To compare the 3 isoforms directly, we first obtained GFP-PP1 expressing cell lines from the Hyman Lab (Dresden, Germany) that were generated using the Bacterial Artificial Chromosome (BAC) strategy[28]. The BAC constructs, randomly integrated into the genome by antibiotic selection, drive the expression of GFP-tagged proteins under the control of their endogenous promoters, which minimizes overexpression artefacts. The PP1α and PP1γ BAC constructs contain the human genes, whereas the PP1β BAC construct contains the mouse gene (100% amino acid and 94% nucleotide identity).

Live cell fluorescence imaging was used to assess the localization of the GFP-tagged PP1 isoforms in HeLa/BAC stable cell lines in which the nucleus was stained with the cell permeable dye Hoechst 33342 (Figure 2a). Their distinct distribution patterns approximated those previously observed^13-15,17^. Given that expression of each tagged isoform is under the control of its endogenous promoter in this system, we were interested to note that GFP-tagged PP1ɑ is expressed at a higher level in cells compared to GFP-tagged PP1β and PP1ɣ. Weaker GFP signals meant that increased exposure times were required to obtain similar fluorescence intensities when imaging the PP1v and PP1 ɣ fusion proteins, as they were not detectable when the same acquisition parameters optimized for imaging GFP-PP1 ɑ were utilized (Supplemental Figure 1a). We observed a similar pattern with Western blot analysis of the tagged fusions in CP/NUC fractions (Supplemental Fig. 1b), and when we estimated the relative abundance of the endogenous isoforms in the U2OS CP:NUC distribution experiment based on the summed intensities (normalized by MW) of the unique peptides identified for each (Figure 1b; Supplemental Data File 1).

**Fig. 2.**
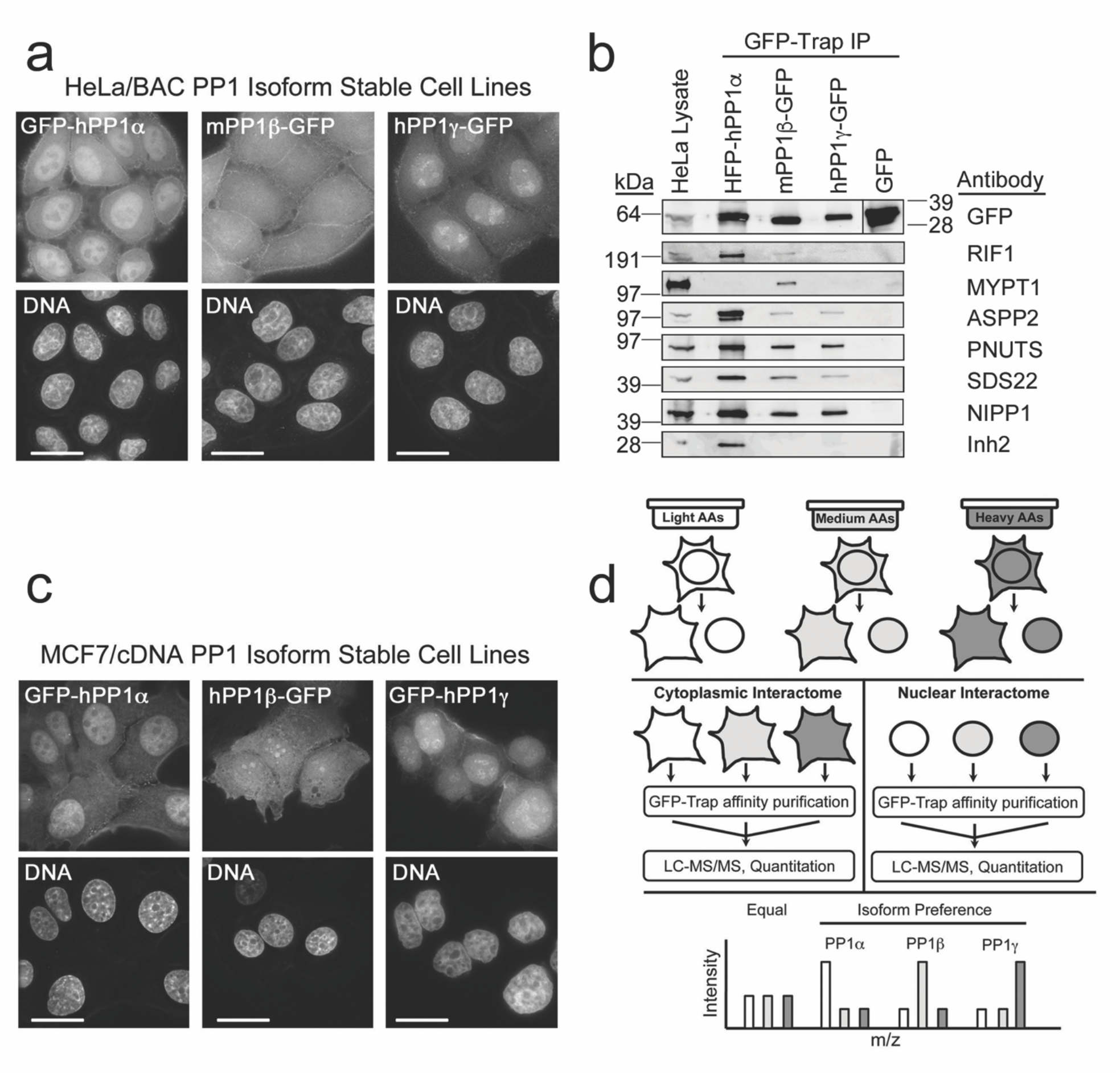
Stable cell lines facilitate the direct comparison of GFP-tagged PP1 isoforms. **(a)** Localization of GFP-tagged PP1 isoforms in live HeLa/BAC cell lines (DNA stained with Hoechst 33342). **(b)** Western blot analysis of GFP-Trap pulldowns of GFP-tagged PP1 isoforms from HeLa/BAC whole cell extracts, using antibodies that recognize a panel of known PP1 regulatory subunits. **(c)** Localization of GFP-tagged PP1 isoforms in live MCF7 stable cell lines (DNA stained with Hoechst 33342). **(d)** Strategy utilized to directly compare the cytoplasmic (CP) and nuclear (NUC) interactomes of GFP-tagged PP1ɑ, β and ɣ stably expressed in HeLa/BAC and MCF7 cell lines. All scale bars are 5 µm.

Association of the GFP-tagged PP1 isoforms with endogenous known regulatory subunits in the HeLa/BAC cell lines was confirmed by AP/Western blot analysis (Figure 2b). Of the 7 regulatory subunits, none were detected in pulldowns of GFP alone (stably overexpressed in HeLa cells by random integration of the pEGFP expression plasmid). As anticipated, the evolutionarily conserved interactors SDS22, NIPP1 and PNUTS were detected in pulldowns of all three PP1 isoforms, whereas the myosin phosphatase subunit MYPT1 showed its expected preference for PP1β.

### Direct comparison of PP1 isoform interactomes identifies a wide range of regulatory proteins and confirms both overlapping and distinct associations

To directly compare the holoenzyme complexes that underlie the subcellular distributions of the three PP1_cat_ isoforms in a non-biased manner, HeLa/BAC cell lines were differentially labeled with Light (PP1ɑ), Medium (PP1β) or Heavy (PP1ɣ) SILAC media as shown in Figure 2d. To reduce sample complexity and increase interactome coverage, cells were fractionated to carry out separate CP and NUC pulldowns. The full datasets (analyzed both individually and concatenated) are provided in Supplemental Data File 2. The same experiment was carried out using MCF7 cell lines established by random integration of pEGFP (CMV promoter) plasmids driving expression of GFP-tagged hPP1ɑ, hPP1β and hPP1ɣ (Figure 2a). Although the expression level of GFP-PP1ɑ is similar to that in the HeLa/BAC lines, the MCF7 cell lines express higher levels of GFP-PP1β and ɣ. The full datasets (analyzed both individually and concatenated) are provided in Supplemental Data File 2.

Comparison of the relative abundance of L, M and H GFP peptides that were detected confirmed that more GFP-PP1ɑ was captured (∼2-4-fold higher than the amounts of GFP-PP1β and ɣ in the MCF7 experiment and >6-fold higher in the HeLa/BAC experiment). This necessarily increases the sensitivity of detection of PP1ɑ-specific complexes, but is representative of physiological conditions as the ɑ isoform does represent a higher fraction of the total pool of PP1_cat_ in cells. Isoform preferences were only noted if a particular regulatory subunit was detected only in the L, M or H form, or was enriched >2-fold above or below the GFP SILAC ratio in the direction of a particular isoform.

Currently, the most comprehensive list of PP1 interacting proteins is annotated as Group ID 694 on the HUGO gene nomenclature website (www.genenames.org), with each assigned a “PPP1R” number to indicate that they are a putative PP1 regulatory subunit. It should be noted that not all of these proteins have been confirmed to form functional complexes with PP1 *in vivo*. There are also a handful of published PP1 regulatory proteins that have not yet been added to this list, such as TPRN (Taperin) and RIF1. We used the PPP1R list (plus 5 additional known interactors) to annotate proteins that were identified in the HeLa/BAC and MCF7 PP1_cat_ interactomes (Table 1). A total of 49 were detected, with substantial overlap observed between the 2 datasets (32 enriched in the HeLa/BAC experiment and 44 in the MCF7 experiment).

**Table 1.**
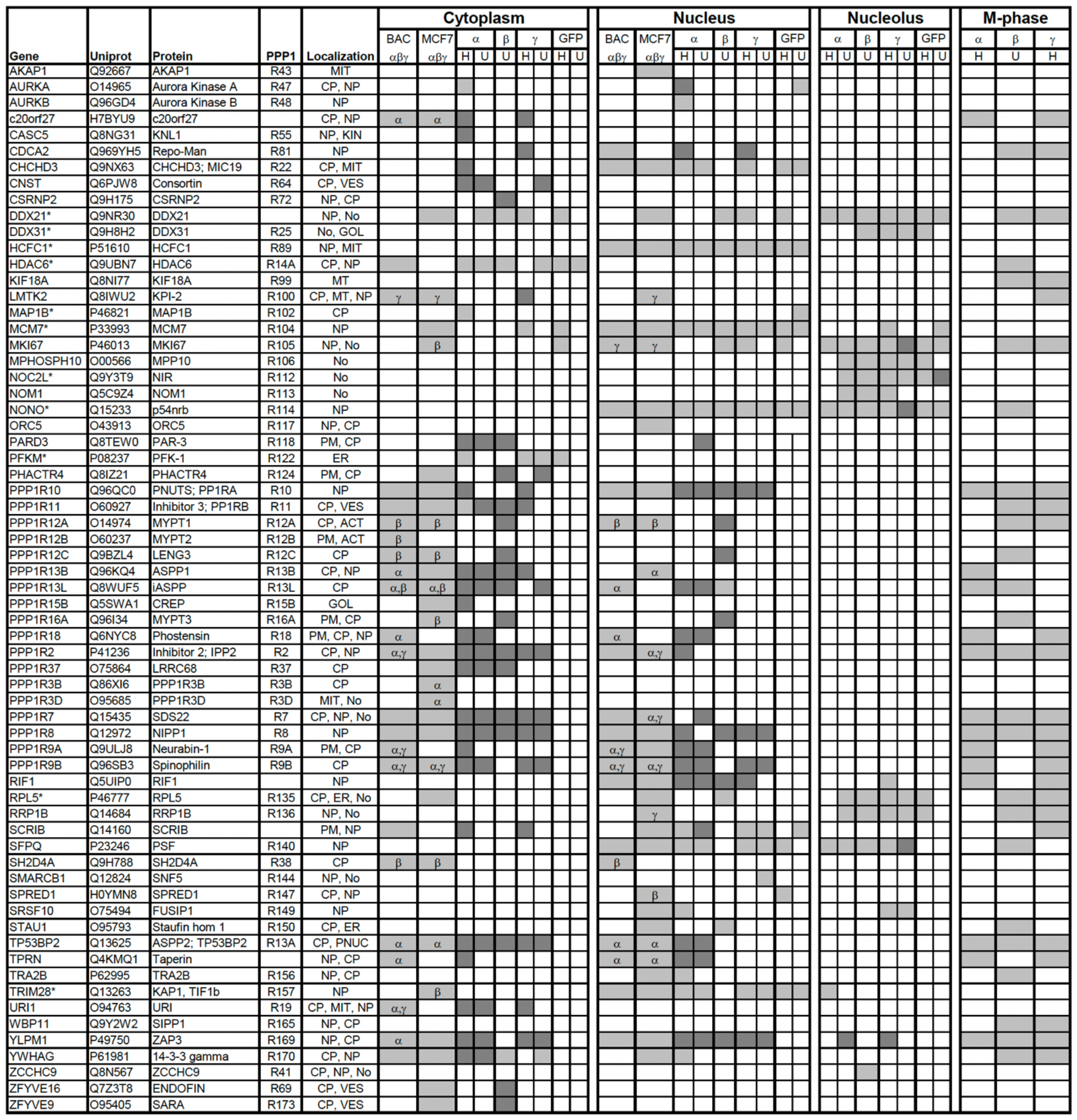
Annotation of known/putative PP1 regulatory proteins detected in the PP1 isoform interactome screens. A light gray box indicates that the protein was detected in an experiment (with any isoform preferences noted for the HeLa/BAC and MCF7 screens). A dark gray box indicates that the protein was enriched >2-fold with PP1 in a focused isoform interactome screen, either in HeLa (H) or U2OS (U) cells. GFP indicates the GFP control datasets. PPP1R numbers are as per HUGO gene nomenclature (group ID 694). Localization information is from the ProteinAtlas online repository (cytoplasm/CP, nucleoplasm/NP, mitochondria/MIT, kinetochore/KIN, vesicles/VES, nucleolus/No, Golgi/GOL, microtubules/MT, plasma membrane/PM, endoplasmic reticulum/ER, actin cytoskeleton/ACT, perinuclear/PNUC). An asterisk (*) indicates that the protein has a high likelihood of binding non-specifically in AP/MS experiments (see Fig. S4).

Some regulatory subunits were only detected in one cell type, which may indicate differences in expression levels. Of the known regulatory subunits identified, most showed the expected isoform preference and compartment specificity. We cloned several for expression in cells as GFP fusions to confirm that their observed localization patterns were consistent in different cell lines. Examples shown in Figure 3c highlight the range of predominantly cytoplasmic to predominantly nuclear localization patterns observed, demonstrating how the steady-state GFP-PP1 localization patterns in interphase cells reflect the weighted summation of their underlying holoenzyme complexes. The importance of relative expression levels in maintaining the balance of PP1 targeting is demonstrated by the disruption observed when a particular regulatory subunit is overexpressed, due to titration of PP1_cat_ away from other holoenzyme complexes (Figure 3d). Although a useful technique for confirming their interaction *in vivo*, this necessarily complicates the analysis of functional experiments based on overexpression of the regulatory subunit (i.e. effects may be due to recruitment of PP1 activity away from other complexes/pathways).

**Fig. 3.**
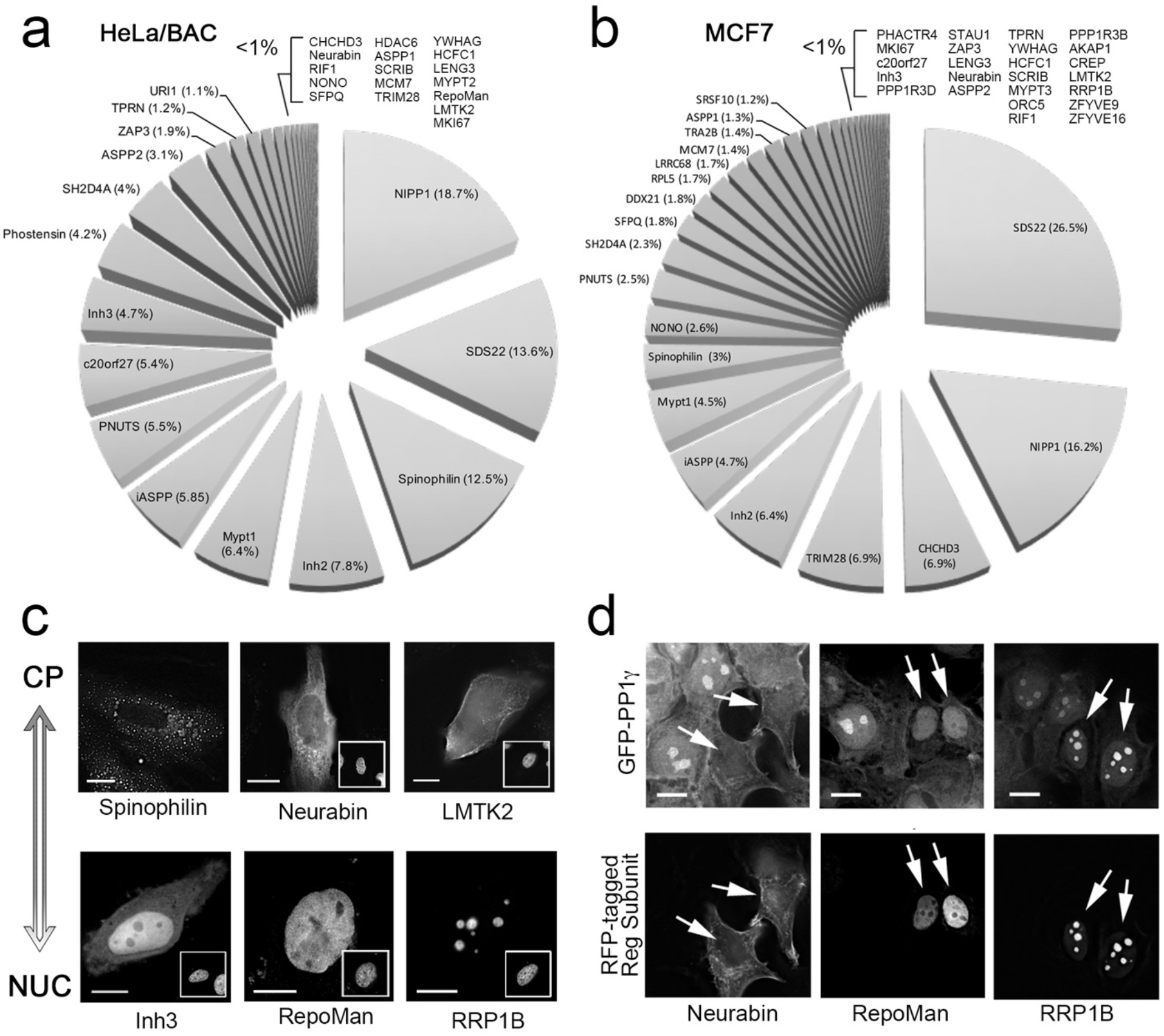
Evolutionarily conserved regulatory proteins represent a large fraction of the PP1c interactome. **(a)** Pie chart summarizing the relative abundance of the known PP1 regulatory proteins identified in the concatenated HeLa/BAC datasets as a fraction of the total abundance of the 32 that were detected. **(b)** Same pie chart analysis for the 43 known PP1 regulatory proteins identified in the concatenated MCF7 datasets. **(c)** Range of subcellular localization patterns for a selection of GFP-tagged regulatory proteins expressed in live HeLa cells (inset: Hoechst-stained nuclei). **(D)** Overexpression of a single red fluorescent protein (RFP)-tagged regulatory protein overrides the steady-state distribution pattern of GFP-PP1ɣ stably expressed in MCF7 cells (arrows). Scale bars are 5 µm.

### Evolutionarily conserved inhibitory subunits represent a large fraction of the holoenzyme complexes captured

Comparison of the peptide intensities (summed and normalized by MW) of the regulatory proteins that co-precipitate with the catalytic subunit provides an indication of their relative abundances in the pulldown. The distribution pie charts in Figure 3a-b plot the relative abundance of each identified regulatory protein as a fraction of the total abundance of all regulatory proteins captured in each experiment (with all 3 PP1 isoforms, and from both the cytoplasmic and nuclear extracts). For both cell lines, the abundant and evolutionarily conserved regulatory proteins SDS22 and NIPP1 represent a significant fraction of the total pool of PP1 holoenzyme complexes captured/identified. Other well-represented regulatory proteins include Inhibitor-2 (Inh2), Spinophilin, iASPP and PNUTS.

It must be stressed that this type of analysis cannot provide absolute stoichiometry for complexes. One reason is variability in the tryptic patterns and number of identifiable peptides for each protein. Also, it is almost certain that we have not identified all of the regulatory subunits with which PP1 is complexed in the cell. This will be due to issues such as detection thresholds (they may represent a very small fraction of the total complexes), affinity (the complex may be disrupted during the AP protocol), cell cycle-specific interactions[29], and the fact that some are not yet on the list of 186 validated interactors for which we screened. For example, having validated the novel protein c20orf27 as a bona fide PP1 interactor over the course of this study (Figure 7), we discovered that it represents a significant fraction (>5%) of the total pool of PP1_cat_ regulatory proteins captured from the HeLa/BAC cell lines. Lastly, some of these regulatory proteins (e.g. SDS22, Inh2) are known to associate with dimeric PP1 complexes to form trimeric holoenzymes, which could account, in part, for their increased representation in the datasets[7].

### Independent fractionation-based AP/MS experiments build a more comprehensive map of the intracellular distribution of individual PP1 isoforms

We next carried out individual mapping of PP1 isoform interactomes using our previously described HeLa GFP-hPP1α and GFP-hPP1γ[15] and U2OS GFP-hPP1α, hPP1β-GFP, and GFP-hPP1γ[17] cell lines. These express higher levels of the β and γ isoforms, like the MCF7 cell lines, and are more amenable to fractionation. Imaging and Western blot analysis confirmed that their subcellular distributions approximate that of their endogenous counterparts (Figure 4a; Supplemental Figure 2).

**Fig. 4.**
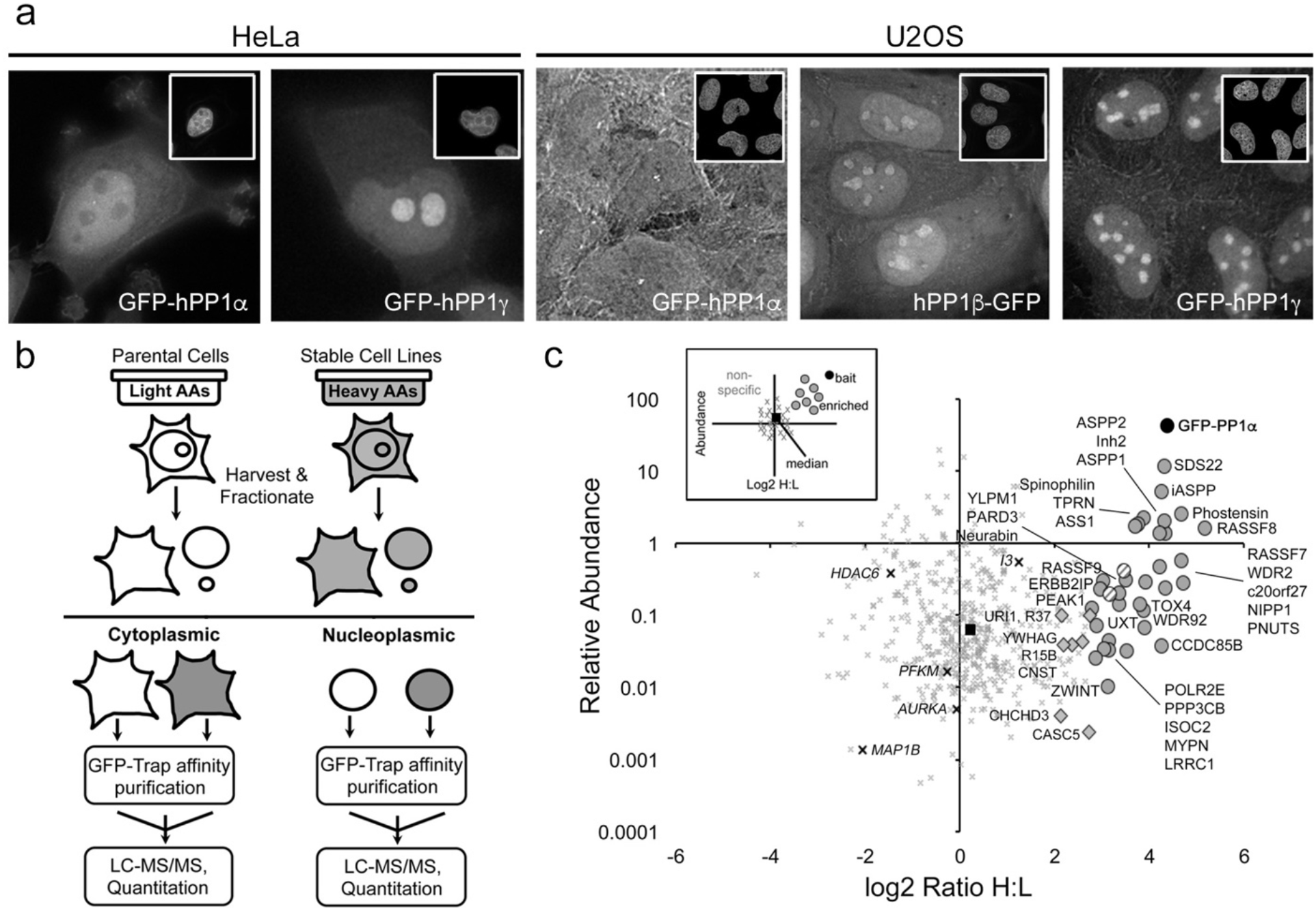
Independent mapping of PP1 isoform interactomes yields high confidence datasets. **(a)** GFP-tagged PP1 isoforms show the expected localization patterns when stably expressed (by random integration of pEGFP expression plasmids) in HeLa and U2OS cells. Hoechst-stained nuclei shown as insets and scale bars are 5 µm. **(b)** Quantitative AP/MS strategy used to independently map cytoplasmic, nucleoplasmic and nucleolar interactomes for each GFP-tagged PP1 isoform in the HeLa and U2OS stable cell lines. GFP-Trap pulldowns from equivalent total protein amounts of parental (L) and PP1 isoform (H) extracts were combined for analysis for each compartmental experiment. **(c)** HeLa cytoplasmic PP1ɑ dataset, plotted as relative abundance (summed and normalized peptide intensities) vs. log2 Ratio H:L. As depicted in the inset diagram, contaminant proteins captured non-specifically in both pulldowns cluster around a median value close to 0 (black square), while GFP-PP1ɑ is highly enriched (black circle). Proteins with Significance B values < 0.05 are indicated (gray circles), as are known regulatory proteins that were enriched > 2-fold (gray diamonds). A black x marks known PP1 regulatory proteins (in italics) that were found at or below the cluster of contaminants, indicating that they likely bound non-specifically in this case. Hashed circles indicate proteins that were enriched in the control GFP dataset. Graphs for all other PP1 isoform datasets are in Fig S3.

To optimally highlight bona fide PP1 isoform interactors above the background of proteins that bind non-specifically to the beads, we differentially labeled parental (L) and GFP-PP1 isoform expressing cells (H) with SILAC media (Figure 4b). Cells were harvested and fractionated, and equivalent total protein amounts of cytoplasmic and nucleoplasmic extracts incubated with the GFP-Trap affinity matrix. In a separate set of control experiments, we compared cell lines stably expressing the GFP tag alone to their respective parental cell line. This was done to highlight contaminant proteins such as chaperones that can be upregulated in unpredictable amounts with exogenous protein overexpression in general, and thus risk being mistaken as enriched in pulldowns of tagged proteins[30]. We identified both cell- and fraction-specific contaminants that were enriched in the GFP control pulldowns (Supplemental Data File 7), and these were flagged in our GFP-PP1 isoform datasets. Examples include chaperones such as HSPA5 and HSPA9, and the short-chain alcohol dehydrogenase DHRS2 that was enriched in nearly all of the GFP pulldowns carried out in the U2OS cell lines. Control and PP1 spatial interactome datasets are provided as supplemental data (Supplemental Data Files 2-5).

To visualize the results of the PP1 interactome datasets, we plotted relative abundance (summed peptide intensity normalized by molecular weight) vs. log2 Ratio H:L (Figure 4c; Supplemental Figure 3) for each protein identified. Contaminants cluster around a Ratio H:L of 1:1 (log2 H:L Ratio 0), as they bind equally under both conditions, while proteins enriched specifically with PP1 have H:L Ratios > 1. We first applied a more stringent Significance B analysis to calculate p values, using the Perseus module of MaxQuant. This takes abundance as well as ratio into account and assigns higher confidence to ratios based on larger numbers of peptide identifications (given that proteins with fewer peptide identifications will show higher ratio variability). For the GFP-PP1α cytoplasmic interactome dataset shown in Figure 4c, 33 proteins (14 of which are known PP1 regulatory proteins) were deemed to be significantly enriched (gray circles). One caveat of this approach is that Significance B values are calculated with respect to the mean ratio. Although ideal for whole proteome comparisons in which the majority of factors do not change while small subsets are enriched or reduced, the mean in AP/MS experiments usually skews toward a higher ratio because there is a cluster at 1:1 (bead contaminants), a few proteins below 1:1 (typically environmental contaminants such as keratins) and then a larger subset with higher ratios that represent the enriched bait protein and interactors. Two-fold enrichment above the median H:L Ratio is thus a strong indicator of a genuine hit, which in this particular dataset expands the list to 98 proteins, including 7 additional known PP1 regulatory subunits (Figure 4c; gray diamonds). We routinely apply both analyses (Significance B values and 2-fold enrichment threshold) to SILAC AP/MS datasets to balance out these differences and to confer different levels of confidence in the results.

Anticipating that some interactors may be close to the 2-fold threshold and occasionally fall below it, we next annotated other known regulatory proteins identified in the screen. Inh3 fell just below the threshold, and is likely to be a genuine interactor in this case. Four other regulatory proteins (HDAC6, PFKM, AURKA and MAP1B) were detected but fell significantly below the threshold, suggesting non-specific binding in this experiment (Figure 4c; black x). We had noted this in other experiments, and set out to assess which of the regulatory proteins in Table 1 are more likely to appear as background contaminants in pulldown experiments. Bearing in mind that regulatory proteins detected in the HeLa/BAC and MCF7 experiments that associate with all 3 PP1 isoforms cannot readily be distinguished from contaminants (as they fall in the same 1:1:1 ratio cluster as proteins that bind non-specifically), we first flagged those that were detected in a control GFP dataset and were not enriched in at least one individual PP1 isoform pulldown. We then looked at the number of control experiments in the CRAPome contaminants repository[31] in which they were detected. Assuming that high abundance sticky proteins are more likely to be detected in control datasets, we next calculated their average abundance in HeLa, MCF7 and U2OS cells, based on values in the ProteomicsDB online repository that were approximated from deposited whole proteome datasets using the iBAQ and top3 intensity approaches[32]. This highlighted a handful of PP1 regulatory subunits that are relatively abundant and commonly observed in control pulldowns (Supplemental Figure 4; * in Table 1). While this does not preclude them being bona fida PP1 regulatory proteins (e.g. a mitosis-specific DDX21/PP1 complex has been validated[33, p. 21]), they should be treated with caution when identified in interactome experiments and rigorously validated using complementary approaches. Because our quantitative approach allows us to apply a stringent confidence threshold to accept whether or not a regulatory subunit has been specifically enriched in a particular experiment, we chose to omit these sub-threshold interactors when assessing the distribution of PP1 between the total pool of holoenzyme partners identified.

This same analysis was applied to all of the PP1 isoform spatial interactome datasets (Supplemental Figure 3) and the results added to Table 1. An additional 16 known regulatory proteins were identified in the HeLa and U2OS single isoform interactomes, bringing the total annotated to 65. As in the HeLa/BAC and MCF experiments, the majority showed the expected isoform preference. For example, the MYPT family members (MYPT1, MYPT2, MYPT3 and LENG3) were only found in the PP1β datasets, while TPRN was limited to the PP1α datasets.

Although we originally intended to include nucleolar interactomes, the reliable identification of interactors proved challenging due to the high background (i.e. non-specific binding of abundant DNA and RNA binding proteins in the extracts). We had previously used this approach to identify RRP1B as a novel PP1ɣ nucleolar regulatory protein, mining the near threshold hits for those that contained putative RVXF motifs. For these new datasets, we annotated all of the known regulatory proteins that were identified (Table 1). Several that have been shown to accumulate in nucleoli (RRP1B, MPP10, NOM1, MKI67) were identified in our nucleolar PP1β and ɣ datasets, but detection of most of them in the control nucleolar dataset suggests that they can also exhibit non-specific binding under these AP conditions, which makes it difficult to detect them reliably above threshold. We will continue to optimize alternate approaches to monitor the sub-nucleolar distribution of PP1.

### Distribution of PP1 isoforms in holoenzyme complexes changes at the entry to mitosis

In general, there was good agreement between the interactomes mapped for PP1ɑ and PP1ɣ in the two cell types, although more interactors were identified in the HeLa cell lines (which express slightly higher levels of the fusion proteins). The bubble graphs in Figure 5 summarize the steady-state distribution of these isoforms in the HeLa lines, and PP1β in the U2OS cell line. For each known regulatory protein that was enriched >2-fold with PP1, its relative abundance in the experiment was plotted vs. its log2 Ratio H:L. The size of the bubble reflects the fraction of the total pool of enriched regulatory proteins that it represents (based on the summed, normalized intensity of H peptides identified for each). This gives a rough approximation of the steady-state distribution of PP1 between the identified holoenzyme complexes. As observed for the HeLa/BAC and MCF7 cell lines, SDS22, NIPP1 and Inh2 represents a large fraction of the complexes identified for all 3 isoforms. Overlap was observed (e.g. ASPP1 and ASPP2 captured with all 3 isoforms), along with isoform-specific complexes (e.g. MYPT1 enriched only in the PP1β dataset; Spinophilin in the PP1ɑ and PP1ɣ datasets).

**Fig. 5.**
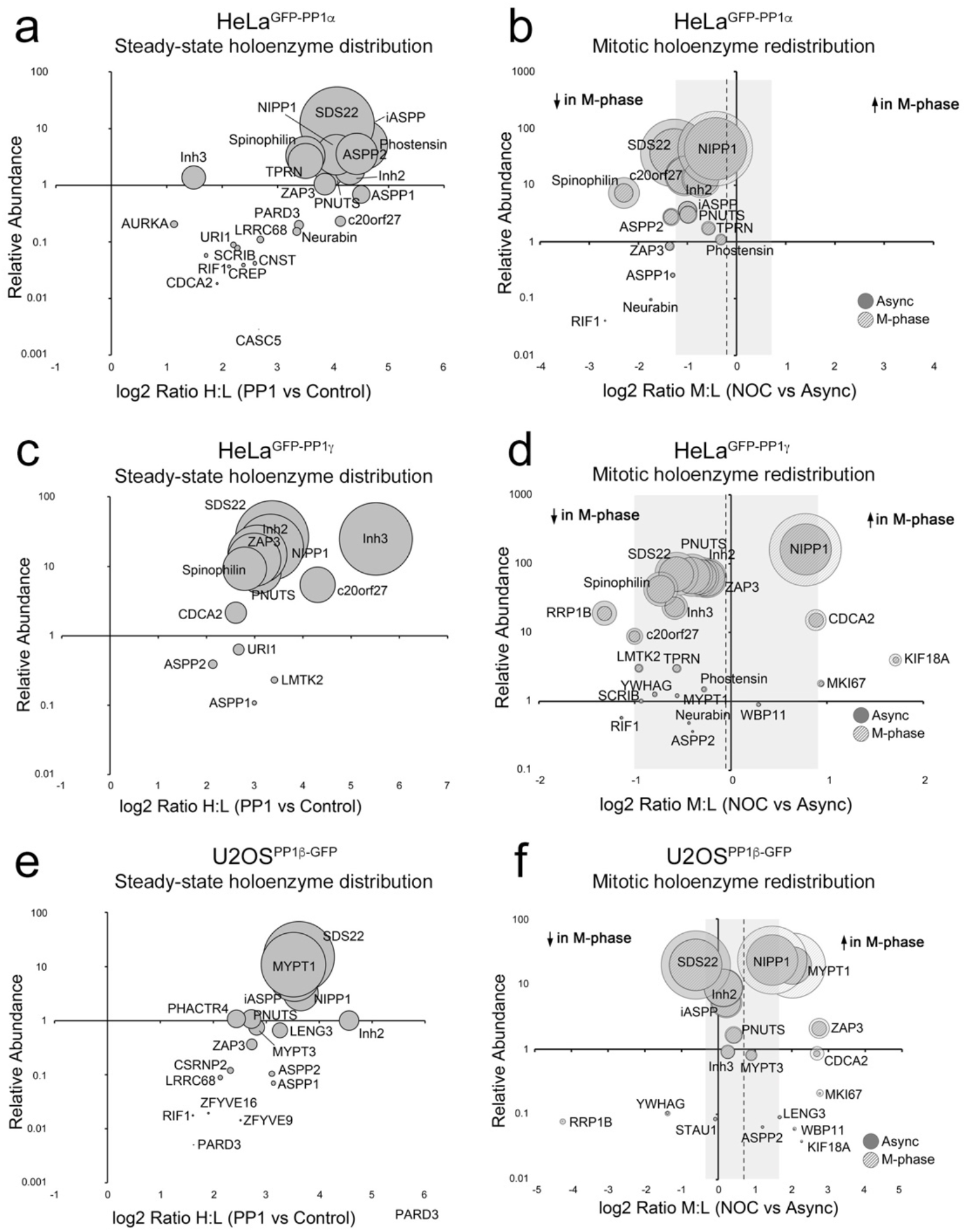
The subcellular distribution of PP1 in known holoenzyme complexes is dynamic. Bubble graphs were used to visualize the subcellular distribution of the 3 PP1 isoforms in holoenzyme complexes under steady-state conditions by plotting relative abundance vs. log2 ratio H:L (PP1 pulldown:control) for all known regulatory proteins enriched >2-fold in pulldowns of HeLa GFP-PP1ɑ **(a)**, HeLa GFP-PP1ɣ **(c)** and U2OS PP1β-GFP **(e)**. The size of the bubble reflects the fraction of the total pool of enriched regulatory proteins that each represents (based on the summed, normalized intensity of Heavy peptides identified). Pulldowns of each isoform were also compared for whole cell extracts from cells arrested at G2/M by treatment with nocodazole (NOC) vs. cells left growing asynchronously (Async). Results are plotted in bubble graphs for HeLa GFP-PP1ɑ **(b)**, HeLa GFP-PP1ɣ **(d)** and U2OS PP1β-GFP **(f)**. In this case, log2 Ratios M:L >2-fold above or below (gray box) the GFP ratio (dashed line) indicate increased enrichment with PP1 under steady-state (L) or M-phase arrested (M) conditions. The size of the gray bubbles reflects the fraction of the total pool of regulatory proteins that each represents in Async cells (summed, normalized intensity of L peptides) and the size of the hashed bubbles reflects the fraction of the total pool of regulatory proteins that each represents in M-phase cells (summed, normalized intensity of M peptides).

This type of analysis also facilitates the identification of changes in the distribution of PP1 that occur in response to a cellular perturbation or at a specific stage of the cell cycle. As an example, we compared the regulatory proteins enriched with the three isoforms in asynchronous cells vs. cells arrested at G2/M. This was done by mapping whole cell interactomes for cells labeled with Light SILAC media (untreated; Async) or Heavy SILAC media (Nocodazole-treated; NOC). The full datasets are provided as Supplemental Data File 6. In the bubble graphs shown in Figure 5, the relative abundance of each regulatory protein identified is plotted vs. the log2 Ratio of NOC:Async. The size of the gray bubbles reflects the fraction of the total pool of regulatory proteins that each represents in Async cells (based on the summed, normalized intensity of L peptides), while the size of the hashed bubbles reflects the fraction of the total pool of regulatory proteins that it represents in M-phase cells (based on the summed, normalized intensity of H peptides). The dashed line is the log2 Ratio H:L for GFP, which marks where an equivalent amount of the isoform was captured in both experiments. Regulatory proteins that fall within the gray zone (< or > 1 log around the dashed line) are considered equally enriched with PP1 under both conditions, while those that fall outside it are considered to be enriched >2-fold in a particular condition.

The most dramatic changes were observed for PP1 ɣ and PP1 β which show increased association with regulatory proteins such as CDCA2/RepoMan, KIF18A and MKI67 that are known to play key roles in mitosis, and decreased association with the nucleolar regulatory protein RRP1B (Figure 5d and 5f). Similar results were observed when cells were arrested at M-phase using Taxol or S-trityl-l-cysteine/STLC (Supplemental Data File 6). Interestingly, PP1ɑ shows no increase in mitotic associations, but rather a net decrease in the overall abundance of regulatory proteins captured. This isoform has been shown to be regulated by inhibitory phosphorylation (Thr320) in mitosis, which may reduce the need for association with certain inhibitory regulatory subunits at this stage. A direct comparison of PP1ɣ (H) vs. PP1ɑ (L) complexes in NOC-arrested cells (Supplemental Data File 6) did show comparable enrichment of several overlapping binding partners, including SDS22, NIPP1 and Ihn2. The expected selective enrichment of RepoMan, KIF18A and MKI67 with PP1ɣ was again observed in this experiment, as was selective enrichment of the ASPP1/2 and iASPP proteins with PP1ɑ.

### Direct vs. indirect interaction with PP1

The identification of a protein in the PP1 interactome, or conversely, detection of PP1 in that protein’s interactome, suggests that they are in a complex but does not imply direct interaction. A key first step in defining their relationship is to determine whether or not the protein possesses any known PP1 binding motifs. As previously noted, the majority of PP1 regulatory subunits (and some substrates) contain a variant of the “RVxF” motif, which can bind to a hydrophobic groove on the catalytic subunit. Although more stringent consensus sequences have been developed for in silico screening[22][34], a good starting point when assessing on a single protein basis is the five-residue consensus motif that covers ∼90% of all known RVxF variants in PP1 binding proteins: [RK]-X(0,1)-[VI]-{P}-[FW])[35]. The caveat is that this motif occurs randomly in around a quarter of all proteins[36].

If the protein does not contain a putative RVxF motif, interaction with PP1 is likely indirect and further analysis of the dataset may highlight a known regulatory subunit that mediates their association. As an example, STRING analysis (http://www.string-db.org)[37] confirmed that the PP1ɑ datasets contain clusters of complexes that point to roles in centrosome regulation, multi-subunit complex assembly and mitochondrial function. This isoform is unique in that it is the only one that displays a detectable accumulation at centrosomes (Figure 6a), and the most enriched centrosomal regulatory protein that was detected with this isoform was ASPP2 (Figure 6b). A closer look at the centrosomal STRING cluster (Fig. 6d) suggested that the RASS family proteins that were detected in our high confidence datasests (RASSF7, RASSF8 and RASSF9) but lack obvious RVxF motifs associate with PP1 indirectly. Consistent with their identification in ASPP1/2 interactome studies[38] [39], all three localize to centrosomes (Figure 6c) and co-immunoprecipitate ASPP2 (Figure 6e). Two other high confidence hits that we identified in our PP1ɑ datasets, CCDC85B/C, are also part of this interaction network (Figure 6c; Table 2).

**Fig. 6.**
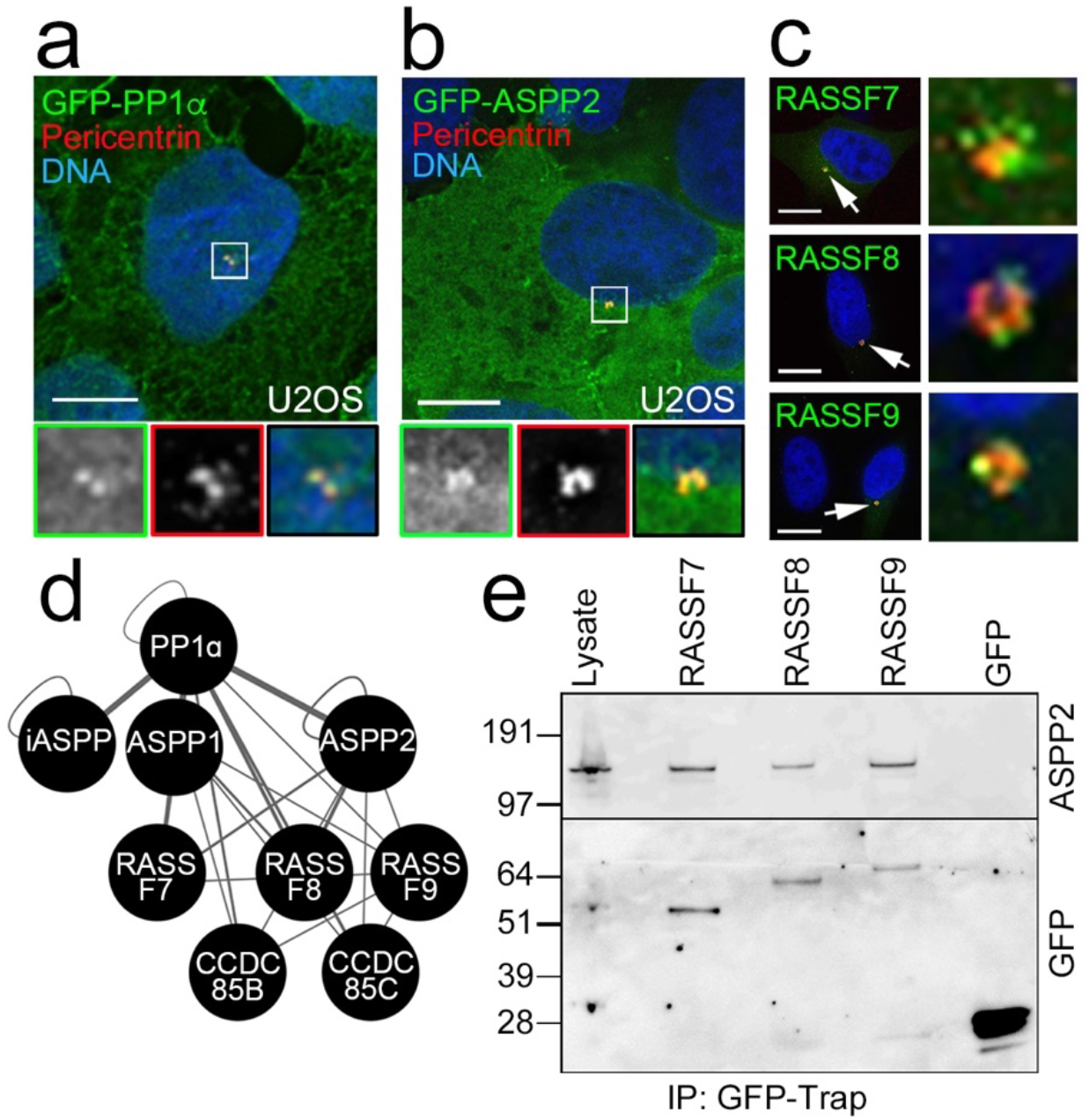
RASSF7/8/9 associate indirectly with PP1ɑ. **(a)** GFP-PP1ɑ shows a visible accumulation at centrosomes, as confirmed here by fixing and counter-staining GFP-PP1ɑ (green) expressing HeLa cells for the centrosomal marker pericentrin (red). DNA was stained with Hoechst (blue). **(b)** The PP1 regulatory protein ASPP2 shows a similar accumulation at centrosomes when expressed in HeLa cells as a GFP fusion protein. **(c)** Expression of GFP-tagged RASSF7/8/9 (green) in HeLa cells confirmed that all 3 proteins accumulate at centrosomes marked by anti-pericentrin (red). (**d)** Proteomic studies have identified associations between a cluster of proteins that were found to be enriched in our PP1ɑ cytoplasmic interactome: ASPP1/2, RASSF7/8/9 and CCDC85B/C. **(e)** Western blot analysis of GFP-Trap pulldowns of GFP-tagged RASSF7, 8 and 9 from HeLa whole cell lysates confirmed co-precipitation of endogenous ASPP2 with all three fusion proteins. The lysate lane is an aliquot of the GFP-RASSF7 lysate prior to pulldown. Scale bars are 5 µm.

**Table 2.**
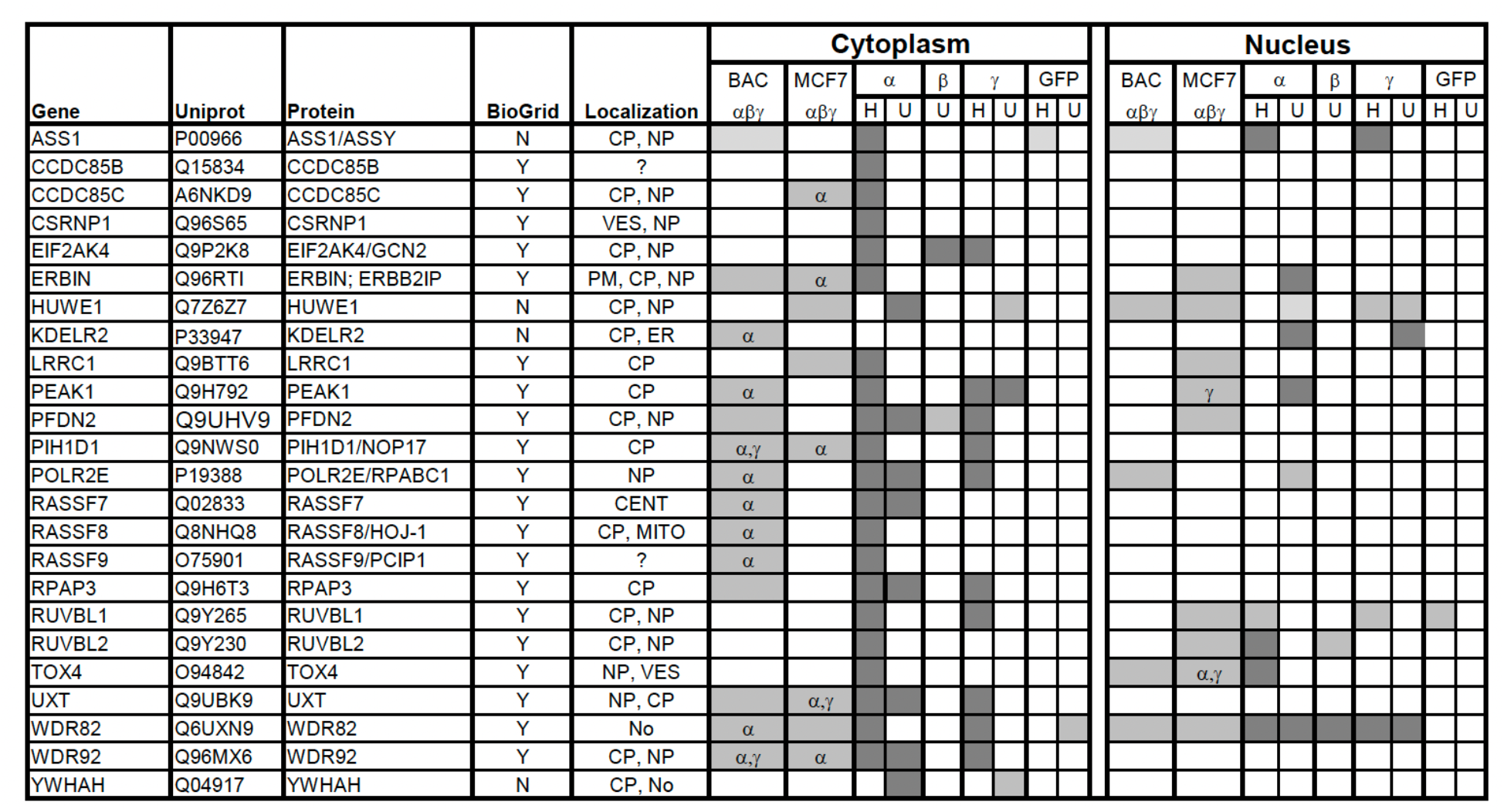
Other high-confidence interactors detected in the PP1 isoform interactome screens. This table lists high confidence hits, most identified in more than one interactome screen. A light gray box indicates that the protein was detected in an experiment (with any isoform preferences noted for the HeLa/BAC and MCF7 screens). A dark gray box indicates that the protein was significantly enriched with PP1 in a focused isoform interactome screen, either in HeLa (H) or U2OS (U) cells. GFP indicates the GFP control datasets. Localization information is from the ProteinAtlas online repository as per Table 1. Also indicated is whether or not the protein has been annotated as a PP1 interactor in the BioGrid database (Y = Yes; N = No).

We flagged an additional 19 high confidence interactors that were identified in more than one PP1 dataset and enriched in at least one of them (Table 2). Many of these have since been deposited in the BioGrid database as putative PP1 interactors annotated in large-scale interactome screens. Most do not contain a putative RVXF motif, meaning that their association with PP1 is likely indirect (i.e. mediated by a regulatory subunit). One example is the PP1 - PNUTS – TOX4 – WDR82 complex[40].

A high confidence hit that did contain a putative RVxF motif was c20orf27 (Figure 7a). Little was known about the protein, and so we first set out to determine if it directly binds PP1, and if that binding is mediated by the KVGF consensus sequence (aa 54-57). Direct interaction with PP1 was confirmed using a bimolecular fluorescence complementation (BiFC) assay, in which the binding partners are fused to the N- and C-termini of YFP (Yellow Fluorescent Protein) and the appearance of yellow fluorescence in the cell indicates direct binding and re-formation of the active fluorophore (Figure 7b). No fluorescence was observed when the “KAGA” mutant c20orf27 was co-expressed with PP1, confirming that their association is mediated by the RVxF motif. Similar results were observed using a fluorescence two-hybrid approach in which accumulation of a GFP-tagged protein at an exogenous gene locus at which mCh-LacR-PP1 is tethered is a readout of their association (Figure 7c). We employed our quantitative AP/MS technique to map the whole cell interactome of GFP-tagged c20orf27 transiently overexpressed in HeLa and MCF7 cells. The top two hits (>2-fold enriched in both experiments) were PP1 and APPBP2. We then repeated the HeLa cell experiment, fractionating cells for separate CP and NUC analysis, to improve our interactome coverage. PP1 and APPBP2 were again identified as top hits, along with other proteins that have been linked to APPBP2 as part of an E3 ubiquitin ligase complex (Figure 7d; Supplemental Data File 8). To rule out the possibility that we are simply capturing c20orf27 that is being targeted for proteasomal degradation by this complex, we demonstrated that the amount of endogenous APPBP2 that co-precipitates with c20orf27 does not change with proteasomal inhibition, nor does GFP-c20orf27 show evidence of a ladder of ubiquitinated forms under this condition (Figure 7e). Future work will assess the contribution of c20orf27-PP1 holoenzyme activity to the functional role of this E3 ubiquitin ligase complex.

**Fig. 7.**
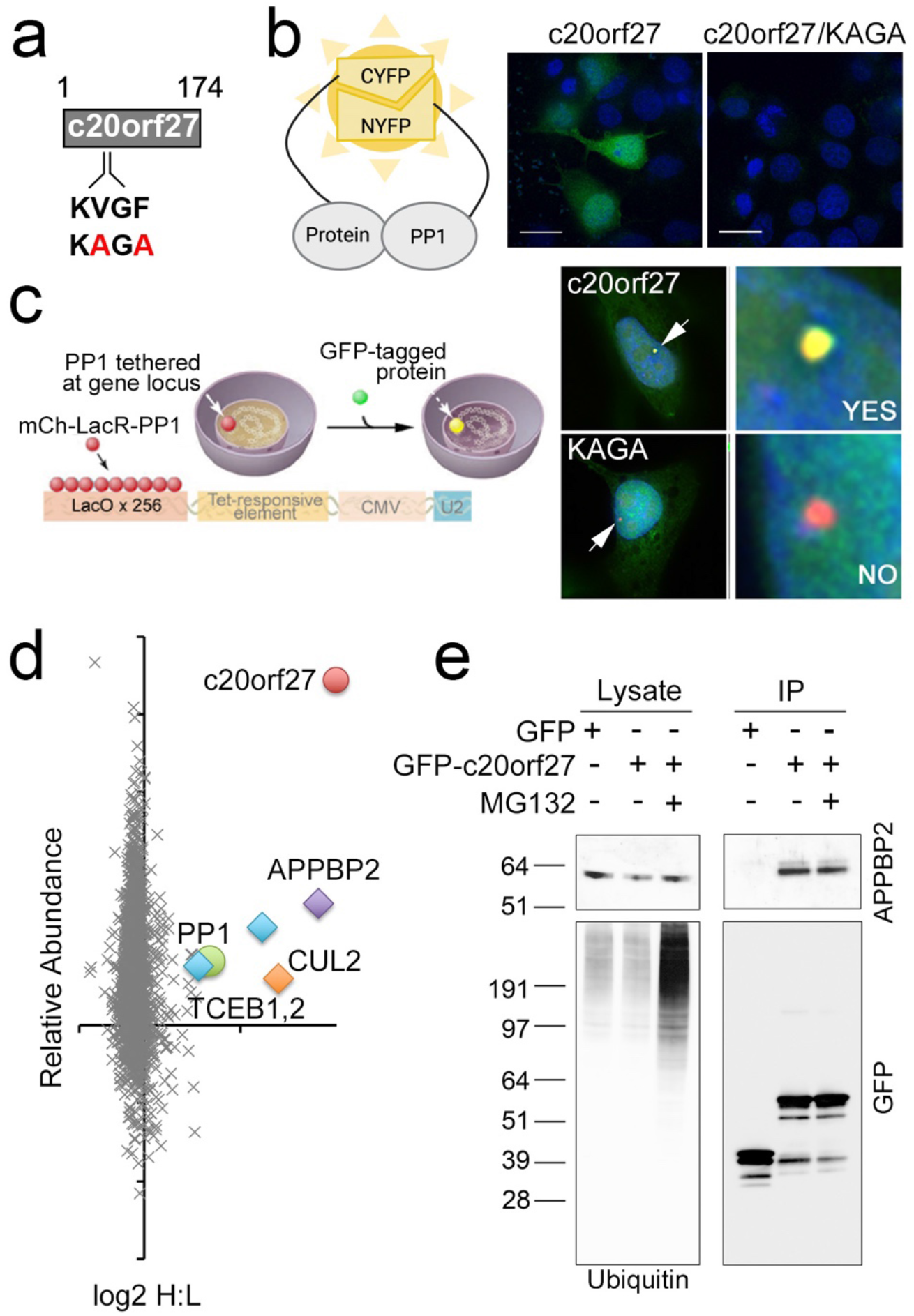
c20orf27 binds PP1 directly. **(a)** Location of the putative RVxF motif in the c20orf27 sequence (aa 54-57), indicating the hydrophobic V and F residues that were mutated to A. **(b)** Bimolecular fluorescence complementation (BiFC) assay confirming that WT but not mutant c20orf27 binds to PP1. The green signal indicates re-formation of functional yellow fluorescent protein (YFP) as the interacting proteins bring its two halves together. **(c)** Fluorescence two-hybrid assay confirms that wild type c20orf27 but not the KAGA mutant associates with PP1 tethered at an exogenous gene locus. **(d)** Quantitative AP/MS dataset for GFP-c20orf27 transiently expressed in HeLa cells, highlighting the high confidence hits. **(e)** Co-precipitation of endogenous APPBP2 with GFP-c20orf27 in the presence and absence of the proteasomal inhibitor MG132. Pulldown of GFP alone is included as a negative control. Scale bars are 5 µm.

## Discussion

In this study we set out to assess the predominant holoenzyme complexes that underlie the distinct subcellular localization patterns observed for the 3 isoforms of the PP1 catalytic subunit. Previous studies have suggested some degree of functional redundancy, but the degree of overlap remains unclear, as does the extent and importance of their unique roles. Comparing the PP1 isoform interactomes in 3 different cultured human cell lines allowed us to identify patterns in their association with specific regulatory proteins and highlight those that represent a significant fraction of the total pool of subcellular PP1 complexes. An interesting observation was the higher level of expression of PP1ɑ relative to the other isoforms in the HeLa/BAC cell lines, in which they are under control of their endogenous promoters. This supports previous suggestions that the ɑ isoform is the most abundant[41]. Observing the same pattern when expression is driven by an exogenous promoter suggests that cells are set up for higher expression of PP1ɑ, for example by expressing certain key regulatory proteins in excess.

Having shown that PP1 is more equally distributed between the cytoplasm and nucleus compared to PP2 and PP5 family members, we incorporated a fractionation step into our interactome mapping workflow. Analyzing cytoplasmic and nuclear extracts separately provides spatial information while also increasing the sensitivity of detection of complexes. Subsequent analysis of concatenated datasets provided whole cell interactomes. Directly comparing relative enrichment of each known/predicted regulatory subunit with the three isoforms enabled us to extrapolate information such as isoform preference and the estimated abundance of a particular holoenzyme complex in the pool of PP1 complexes recovered. Many of the preferences that we observed are in agreement with previously published results, such as the preferential association of TPRN with the alpha isoform[27] and the MYPT family members with the beta isoform[42], [43]. Interactome data deposited in the BioGrid database lists LMTK2 as both a PP1α and PP1γ interactor. The fact that LMTK2 was only detected with PP1γ in our HeLa/BAC screen, however, despite capturing nearly 3x more of the α isoform on the beads, suggests that it shows a preference for the γ isoform in vivo. For this and other regulatory subunits, the functional significance of preferential association with a particular isoform remains to be determined. An important caveat to assigning isoform specificity is that overexpression of a particular regulatory subunit can override preferences. Our lab previously demonstrated this for the chromatin-associated regulatory subunit RepoMan, which we originally identified via its selective association with stably expressed GFP-PP1ɣ over GFP-PP1ɑ. When overexpressed, RepoMan can recruit all 3 isoforms to chromatin, disrupting their normal subcellular distribution in interphase and mitotic cells[15]. This highlights the strong potential for off-target effects when assessing the functional role of a particular regulatory subunit by overexpression, as it is difficult to separate the direct effect of excess PP1 recruitment to a particular site from the indirect effect of recruiting it away from other substrates. Based on our interactome results, together with results deposited in databases such as BioGrid and the published literature, we chose to avoid the term “isoform-specific” here, and only refer to isoform “preferences”. Table 1 summarizes the total of 65 known regulatory proteins that were detected in our screens, noting any isoform preferences observed in the HeLa/BAC and MCF7 experiments and in which individual isoform pulldowns they were identified (and if they were >2-fold enriched with PP1).

The most striking result from our screens was the consistent finding that a significant fraction of PP1_cat_ associates with the evolutionarily conserved regulatory subunits SDS22, NIPP1, Inhibitor-2 (Inh2) and Inhibitor-3 (Inh3). In the HeLa/BAC and MCF7 screens, these 4 subunits accounted for approximately 45-54% of the total pool (by relative abundance) of regulatory subunits identified. A similar pattern was observed in the individual isoform interactomes, with SDS22 consistently dominating the cytoplasmic pool of interactors and NIPP1 the nuclear pool. It is important to note that although all four of these proteins were initially identified as PP1 inhibitors, that distinction cannot readily be made for phosphatases because it depends on the substrate that is used to assess inhibition. For example, the glycogen regulatory subunit increases the activity of PP1 against glycogen phosphorylase, but inhibits its activity against myosin. Conversely, the myosin regulatory subunit activates the activity of PP1 against myosin while inhibiting its activity against glycogen phosphorylase. The evolutionarily conserved “inhibitory” subunits have been shown to positively regulate PP1 in specific pathways as well as inhibiting its activity against unrelated substrates[44], [45].

That said, the strong representation of SDS22, NIPP1, Inh2 and Inh3 in all 3 PP1 isoform interactomes is consistent with previous studies that suggest that they play key roles either as chaperones in the initial maturation of translated PP1 or as PP1 “sinks” that bind and inhibit activity of the free catalytic subunit in the cell until it is recruited by other regulatory subunits as needed[46], [47]. SDS22 binds PP1 via a leucine-rich repeat region rather than an RVxF motif, which allows it to form trimeric complexes as the “3rd subunit” of multiple PP1 holoenzymes[48]. An inhibitory complex of SDS22-PP1-Inh3, for example, has been described in both yeast[49] and mammalian cells[47]. One suggested role is to limit the association of SDS22 with kinetochore-bound PP1 during mitosis, to prevent premature phosphatase activity at this structure and ensure bipolar mitotic spindle attachment[50]. SDS22 and Inh3 have also been proposed to form a transient complex with the catalytic subunit during PP1 biogenesis, in a process that involves the eventual ubiquitin-independent disassembly of the complex to release free PP1[46].

SDS22 was recently shown to selectively recognize and trap inactive, metal-deficient PP1_cat_[51]. Dissociation of SDS22 is accompanied by binding of the required second metal, which results in an active phosphatase that can be recruited into a holoenzyme complex. This may resolve the conundrum of whether or not SDS22 is a PP1 inhibitor, chaperone or regulatory protein, as it suggests a central role (which incorporates features of each) as a PP1 “sink” or “bank” that holds a pool of catalytic subunits inactive and ready for holoenzyme formation. This would allow cells to respond rapidly to signaling requirements by either synthesizing (or activating) the relevant regulatory subunits. Our finding that SDS22 is the predominant interactor associated with all 3 PP1 isoforms under steady-state conditions, and the shift of a fraction of this pool to mitotic-specific regulators as G2/M, is consistent with an SDS22-mediated PP1 sink being a major mechanism by which PP1 activity is regulated in cells.

Also consistent with this hypothesis is the fact that the regulatory subunits are expressed in excess over the catalytic subunit[7], which would explain how cells can adapt to fluctuations in PP1 levels, including its stable overexpression. In addition to these 4 evolutionarily conserved regulatory proteins (SDS22, NIPP1, Inh2 and Inh3), our PP1 isoform interactome screens identified numerous other regulatory proteins (∼30-40 per experiment), confirming that under steady-state conditions the catalytic subunit is both held in inactive complexes and incorporated into functional holoenzyme complexes throughout the cell. For PP1ɑ, other dominant regulatory proteins include 3 members of the ASPP (apoptosis stimulating proteins of p53) family, the F-actin targeted Phostensin and the nuclear regulatory proteins PNUTS, ZAP3 and TPRN. A somewhat surprising result was the finding that Spinophilin accounts for a relatively large fraction of the PP1ɑ and PP1ɣ interactomes, and was detected in both the cytoplasm and nucleus. This regulatory protein, which has been shown to act as a scaffold that organizes both cytoskeletal and membrane functions, is highly enriched at dendritic spines and thus has primarily been studied with respect to its role in neuronal cells[52]. Its strong representation in our screens suggests key roles in non-neuronal cells, as well. It has also been implicated as a tumor suppressor[53] that can interact with the tumour suppressor ARF[54] and regulate phosphorylation of Rb[55]. It is also known to show a lower affinity for PP1β[56], [57], and was not detected in our PP1β interactomes. Those were dominated by the myosin phosphatase regulatory protein MYPT1, in both the cytoplasm and nucleus. The PP1β nuclear interactome also enriched histones to a greater extent, indicating a stronger association with chromatin. This is consistent with a genome-wide promoter binding study that identified it as the primary promoter-bound isoform[58].

Although we expected our interactome screens to mainly identify the regulatory proteins that are the “hubs” of PP1 signaling in the cell, with limited coverage of additional proteins in each complex, we did observe extensive coverage for certain complexes. One is the R2TP/Prefoldin-like co-chaperone complex (RUVBL1-RUVBL2-RPAP3-PIH1D1). This complex associates with HSP90 to facilitate assembly and stability of a number of critical large protein or protein-RNA complexes, including RNA polymerase II, small nucleolar ribonucleoproteins (snoRNPs) and phosphatidylinositol 3 kinase-related kinases such as mTOR[59]. It was also shown to associate with additional subunits of the prefoldin module (PFDN2, PFDN6, URI1, UXT, PDRG1, POLR2E, WDR92) to form an R2TP/PFDL complex that is believed to play a key role in recruiting proteins for complex assembly[60]. With the exception of PFDN6 and PDRG1, we identified all of these complex members as enriched in our PP1 interactome screens. This suggests that participation in the assembly of large multiprotein (and protein-RNA) complexes is one of the key cellular roles for PP1.

Another interesting observation was that PP1ɑ is the only isoform that accumulates at centrosomes in interphase cells in the three cell lines tested, but the only centrosomal-associated regulatory proteins detected in its interactome were the ASPP1/2 family members. We did not detect other regulatory proteins that have linked PP1 to centrosomal roles, such as the centrosomal kinase NEK2, LRRC67/PPP1R42 or CEP192[61]–[63]. This suggests that the ASPP family may play a more central role in initial recruitment, while the more specialized complexes represent only a small fraction of the total pool of PP1 at centrosomes. Recent structural work demonstrated that ASPP2 binds PP1 via both a canonical RVXF motif and an SH3 domain that engages the C-terminal tail of the catalytic subunit[39], [64]. This enables high affinity and tunable binding, and the ability to discriminate between the PP1_cat_ isoforms (due to their varying C-termini). Although ASPP2 was detected with all three isoforms in our screens, a significantly larger fraction was found in complex with PP1α. In addition, three Ras association domain family (RASSF) scaffold proteins previously linked to ASPP1/2 (RASSF7, F8 and F9) were only identified in our PP1α screens, and we confirmed that all 3 accumulate at centrosomes and associate indirectly with PP1 via ASPP2. Studies in Drosophila linked dASPP, the single homolog of human ASPP1 and ASPP2, to the RASSF8 homolog dRASSF8, and showed that the complex regulates cell-cell adhesion during retinal morphogenesis[65]. The PP1/ASPP/RASSF complex has since been extended to include 2 isoforms of the small coiled coiled domain protein 85 (CCDC85B/C)[65], which we also identified in our PP1ɑ screen.

Functionally, ASPP1/2 have been shown to interact with the centrosomal linker protein C-Nap1, and it was proposed that they play a role in linker reassembly at mitotic exit[66]. Of particular interest, however, is a study suggesting that the ASPP2-PP1ɑ complex may be important in the regulation of the Hippo pathway, which regulates tissue homeostasis[38]. We also identified two other PP1 partners that have been linked to regulation of the Hippo pathway, suggesting that the phosphatase feeds into this signalling cascade at more than one level. These are the regulatory subunit PARD3, which is involved in cell polarity regulation and has been shown to induce TAZ activation by promoting the interaction of PP1 with the LATS1 kinase[67] and PEAK1[68], a protein tyrosine kinase that is associated with the actin cytoskeleton and has been linked to Hippo pathway regulation via eIF5A-PEAK1 signaling-mediated control of YAP1/TAZ expression levels. PEAK1 was one of the commonly enriched interactors observed in our individual isoform interactome screens that contained a putative RVxF motif. Although we confirmed its association with PP1, mutation of the RVxF motif reduced but did not prohibit their association, suggesting either that additional PP1 binding regions strengthen the interaction or that binding is indirect. Future studies are needed to address this in more detail, along with the relative contributions of the various PP1 holoenzyme complexes to Hippo pathway signalling.

Another commonly enriched interactor identified in our screens that we did validate as a novel bona fide PP1 regulatory protein is c20orf27. Little is known about this protein, although a recent study did detect PP1 in its interactome and linked it to the TGFβR-TAK1-NFκB signalling pathway[69]. We first confirmed that its RVxF motif mediates direct association with PP1. Functional clues to its cellular role(s) were then provided by AP/MS screens that not only confirmed co-precipitation of PP1 but also revealed a consistent interaction with APPBP2 (amyloid beta precursor protein binding protein 2), the human homolog of Drosophila PAT1. This protein is primarily known for its upregulation in certain cancers[70], [71], and a possible link to microtubule trafficking of the amyloid precursor protein (APP) and androgen receptor transactivation[72], [73]. Of particular interest to is its identification as a novel VHL-type Elongin BC-box CUL2 binding E3 ubiquitin ligase substrate specifier[74]. In addition to PP1 and APPBP2, the top hits in our c20orf27 interactome were the Cullin protein CUL2 and the ElonginB/C proteins TCEB1/2. The first substrate for this complex was recently identified as the PR domain-containing protein 16 (PRDM16), which plays a key role in biogenesis of beige adipocytes[75]. Targeted depletion of CUL2-APPBP2 in adipocytes activated the PRDM16 pathway to counteract diet-induced obesity and insulin resistance, highlighting the therapeutic potential of this complex with regard to metabolic health. To rule out the possibility that c20orf27 is simply a substrate for this complex, we demonstrated that there is no increase in the amount of APPBP2 associated with c20orf27 when proteasomal function is inhibited. There is also no evidence that c20orf27 is ubiquitinated, either in the presence or absence of proteasome inhibitors.

Although our complex did not include a classic ubiquitin ligase, we did see enrichment of UBR2 (Recognin), which plays a role in a unique branch of the ubiquitin-proteasome system called the “N-end rule pathway”. In this pathway the N-terminal residue of a protein serves as an N-degron, which is marked by various post-translational mechanisms and targeted for degradation[76]. Failures in this pathway have been implicated in the development of human diseases such as Johanson‒Blizzard syndrome and neurodegenerative disorders, and the Alzheimer’s Disease-associated fragment of APP has been identified as an N-end rule pathway substrate[77]. This suggests that the pathway counteracts neurodegeneration, and thus the fact that our novel complex links regulatory phosphatase activity (c20orf27/PP1) to a key N-end pathway factor and an E3 ubiquitin ligase substrate specifier (APPBP2) that also binds APP is of significant therapeutic interest.

In addition to its broad range of regulatory roles in interphase cells, key mitotic-specific roles have also been described for PP1 (see [78] for review) and our data confirm that association with a subset of regulatory complexes increases at G2/M, at least for the PP1β and PP1γ isoforms. Some, such as RepoMan and MKI67, are associated with PP1 throughout the cell cycle and play regulatory roles in interphase, but their expression levels increase prior to mitosis due to essential roles in this process[79]. When cells arrested at G2/M by nocodazole treatment were compared to asynchronous cells, we observed a 2-fold increase in the amount of these proteins associated with PP1γ, and a 5-7-fold increase in the amount associated with PP1β. KIF18A was only detected in our mitotic screens, and showed a 3-4-fold increase in G/M-arrested cells, consistent with its role as a mitotic kinesin[29]. We did not detect any PP1α complexes that were upregulated at G2/M, and in fact there was a general decrease in its association with several regulatory subunits. PP1 has been shown to be regulated in mitosis, at least in part, via inhibitory phosphorylation of a threonine residue (Thr^320^ in PP1α) by the mitotic kinase CDK1[80]. This may indicate that the primary regulatory roles for this isoform are carried out in interphase, and that it remains predominantly inactive throughout mitosis. Alternatively, it may already exist in the complexes that regulate its mitotic role(s), with no need for increased levels of these complexes. Interestingly, although Inhibitor-1 (PPP1R1A) is also believed to play a role in suppressing PP1 activity in early mitosis, we did not detect this regulatory protein in our mitotic screens. Its small size (19 kDa) and tryptic profile may reduce its detectability by MS, which is also suggested by the fact that it was only annotated in 3 cultured cell lines in the ProteomicsDB repository of whole proteome datasets (for comparison, PP1_cat_ was detected in >130 deposited cell line datasets).

The regulatory subunit RIF1, found in our PP1ɑ and PP1ɣ datasets, showed a reduced association with both in M-phase arrested cells. Similarly, RRP1B, which associates preferentially with PP1β and PP1ɣ, also showed a reduced association in M-phase in both datasets. Association of RIF1 and RRP1B with PP1_cat_ was recently shown to be negatively regulated by Aurora B kinase-mediated phosphorylation of a Ser/Thr residue in their RVxF docking motifs (RV[S/T]F) at mitotic entry^⍰⍰^. Our dynamic distribution results independently confirm specific loss of their association with PP1_cat_ at G2/M, supporting the idea that this is a key mechanism employed to regulate PP1 activity throughout the cell cycle. Moving forward, our unperturbed interactome “snapshots” will be useful comparisons for the changes that occur during different cellular conditions (e.g. proliferation vs. differentiation or the epithelial-to-mesenchymal transition) and in response to various cellular perturbations. It will also help us to identify changes that have occurred in various disease states.

## Materials and Methods

### Plasmids and antibodies

The FP-PP1 isoform constructs were previously described[14], [25] and are available through Addgene. For the Fluorescenct-Two-Hybrid experiments, PP1ɣ was subcloned into an mCherry-NLS-LacR[81] expression plasmid. Coding sequences for PP1 interactors were amplified from donor constructs (obtained from Addgene or ThermoFisher) and subcloned into pEGFP-C1/N3 and/or pmCherry-C1/N3 expression plasmids, with in-frame insertion confirmed by DNA sequencing. The coding sequence for PEAK1 was amplified from a donor plasmid kindly provided by the Klemke lab[68] and subcloned into the pEGFP-C1 expression plasmid. In-house PP1 isoform-specific antibodies used for Western blotting were previously described[27]. Details for all primary and secondary antibodies and dyes utilized in this study can be found in Supplemental Table 1.

### Cell culture

HeLa, MCF7, U2OS and HEK293 cells (all from ATCC), were grown in Dulbecco’s modified Eagles’ medium (DMEM) supplemented with 10% fetal calf serum and 100 U/mL penicillin and streptomycin (Wisent Bioproducts Inc, St. Bruno, QC). Stable cell lines were generated as previously described and maintained in media supplemented with G418[82]. The Bacterial Artificial Chromosome (BAC) HeLa cell lines hPPP1CA-NFLAP, mPPP1CB-LAP and hPPP1CC-LAP were a generous gift from Drs. Ina Poser and Anthony Hyman (Dresden, Germany)[28]. They were further subcloned to obtain populations with >95% cells of expressing the fusion proteins. The HeLa cells with stable incorporation of a 256-copy lac operator array used for the fluorescence two-hybrid screens were a gift from Greg Matera (North Carolina, USA)[81].

### Metabolic labeling

Stable isotope labeling with amino acids in cell culture (SILAC) for label-based quantitative MS was carried out as previously described[30]. For double encoding experiments, cells were grown for 7-10 passages in high glucose DMEM containing either L-arginine and L-lysine (Light), or the isotopes L-arginine^13C/15N^ and L-lysine^13C/15N^ (Heavy). For triple encoding experiments, media containing the isotopes L-arginine^13C^ and L-lysine4,4,5,5-D4 (Medium) was also used. SILAC media was prepared by supplementing high glucose DMEM minus Arg/Lys/Leu/Met (AthenaES) with 10% dialyzed FBS (ThermoFisher) and the appropriate amino acids, mixing well and filtering through a 0.22 µm filter (Millipore).

### Preparation of cell extracts

Whole cell extracts were prepared by sonication in ice-cold RIPA buffer (50 mM Tris pH 7.5, 150 mM NaCl, 1% NP-40, 0.5% deoxycholate, protease inhibitors) and cleared by centrifuging at 2800g for 10 min at 4 °C. For mitotic arrest experiments, cells were treated with either 0.1 μg/ml nocodazole, 10 nm Taxol, or 5 μm S-trityl-l-cysteine (STLC) for 18 h before harvesting by mitotic shake-off and preparation of whole cell extracts. All drugs were obtained from Sigma-Aldrich.

Cytoplasmic and nuclear extracts were prepared from subcellular fractions, which were obtained as previously described[83]. Briefly, HeLa cells harvested from 5 × 15 cm dishes were resuspended in 5 ml of ice-cold swelling buffer (10 mM Hepes, pH 7.9, 1.5 mM MgCl2, 10 mM KCl, 0.5 mM DTT, and protease inhibitors) for 5 min, and the cells broken open to release nuclei using a pre-chilled Dounce homogenizer (20 strokes with a tight pestle). Dounced cells were centrifuged at 228 g (1,000 rpm, GH-3.8 rotor; Beckman Coulter GS-6) for 5 min at 4°C to pellet nuclei and other fragments. For U2OS and MCF7 cells, which are less amenable to douncing, harvested cells were resuspended in 4 mL of ice-cold mild detergent buffer (20 mM Tris pH7.4, 10 mM KCl, 3 mM MgCl2, 0.1% NP40, 10% glycerol) for 10 min, followed by centrifugation at 1350g for 10 min at 4 °C to break open cells and release and pellet nuclei and other fragments.

For both methods, the supernatant was retained as the cytoplasmic fraction while the nuclear pellet was resuspended in 3 mL of 0.25 M sucrose/10 mM MgCl2, layered over a 3 mL cushion of 0.35 M sucrose/0.5 mM MgCl2 and centrifuged at 1430g for 5 min at 4 °C. Cytoplasmic extracts were prepared by adding 1 mL of ice-cold 5× RIPA buffer to the 4 mL cytoplasmic fraction, following by a brief sonication and centrifugation at 2800g for 10 min at 4 °C. Nuclear extracts were prepared by resuspending the purified nuclei in 5 mL of 1× RIPA buffer, sonicating, and clearing by centrifugation. For additional fractionation of nuclei into nucleoplasmic and nucleolar fractions, the nuclear pellet was resuspended in 0.35 M sucrose/0.5 mM MgCl2 and sonicated to disrupt nuclei and release nucleoli, which are visible by light microscopy as dense, refractile bodies. The sonicate was layered over a 3 ml cushion of 0.88 M sucrose/0.5 mM MgCl2 and centrifuged at 2800 g for 10 min at 4°C to pellet nucleoli and the supernatant retained as the nucleoplasmic fraction. Nucleoplasmic extracts were prepared by adding 2 ml of ice-cold 5X RIPA buffer and 2 ml of ice-cold dH2O to the 6 ml nucleoplasmic fraction, sonicating and clearing by centrifugation. Nucleolar extracts were prepared by resuspending purified nucleoli in high salt (500 mM) RIPA buffer for sonication and centrifugation. Total protein concentrations were measured using the Pierce BCA Protein Assay Kit (ThermoFisher).

### Affinity purification

GFP-PP1 complexes were captured from cell extracts as previously described[30], with equal amounts of total protein extract for each condition incubated with GFP-Trap_A beads (Chromotek) at 4°C for 1 hour. For pulldowns from nucleolar extracts, the salt concentration was reduced to 250 mM by adding an equal volume of “no salt” (0 mM NaCl) RIPA buffer. Following an initial wash with RIPA buffer, beads from the control and experimental pulldowns (or 3 different PP1 isoform pulldowns) were combined for additional washes (to minimize variability in downstream processing) and bound proteins eluted with 1% SDS. The eluted proteins were reduced and alkylated by treatment with DTT and iodoacetamide, respectively. Sample buffer was added and the proteins resolved by electrophoresis on a NuPAGE 10% BisTris mini gel (Thermo Fisher). The gel was stained using SimplyBlue Safestain (Thermo Fisher) and the entire lane cut into five slices. Each slice was cut into 2 × 2 mm fragments, destained, and digested overnight at 30°C with Trypsin Gold (ThermoFisher).

For direct comparison of cytoplasmic vs. nuclear proteomes, cell equivalent volumes of each extract (differentially encoded) were combined for separation on a NuPAGE 4-12% Bis-Tris gel and the lane cut into 12 slices for in-gel trypsin digestion and MS analysis.

### Mass spectrometry and data analysis

An aliquot of each tryptic digest was analyzed by LC-MS/MS on an Orbitrap Fusion Lumos system (Thermo Scientific) coupled to a Dionex UltiMate 3000 RSLC nano HPLC. The raw files were searched against the UniProt human database using MaxQuant software v1.5.5.1 (http:/www.maxquant.org)[84] and the following criteria: peptide tolerance = 10 ppm, trypsin as the enzyme (two missed cleavages allowed), and carboxyamidomethylation of cysteine as a fixed modification. Variable modifications are oxidation of methionine and N-terminal acetylation. Heavy SILAC labels were Arg6 (R6), Arg10 (R10), Lys4 (K4), and Lys8 (K8). Quantitation of SILAC ratios was based on razor and unique peptides, and the minimum ratio count was 2. The peptide and protein FDR was 0.01.

### Fluorescence microscopy

Cells seeded on No. 1.5 coverslips were fixed for 10 min at room temperature in 3.7% (wt/vol) paraformaldehyde (PFA) in CSK buffer (1mM PIPES pH 6.8, 100mM NaCl, 300mM sucrose, 3mM MgCl2, 2mM EDTA). Following a 10 min permeabilization with 1% Triton X-100 in PBS, nonspecific binding sites were blocked by incubation in PBS with 1% BSA and 0.2% Tween-20. For immunostaining, cells were incubated with the appropriate primary antibody, followed by incubation with the appropriate fluorophore-conjugated secondary antibody. Biotinylated proteins were stained by incubating cells with Alexa Fluor 647-conjugated streptavidin (ThermoFisher). Coverslips were prepared for imaging by mounting in Vectashield liquid mounting media (Vector Labs).

For live imaging, cells were cultured in 35 mm optically clear polymer-bottom µ-dishes (ibidi) and growth medium replaced with Phenol Red-free CO_2_ independent medium (ThermoFisher). If desired, DNA was stained by incubating the cells for 20 min at 37°C in medium containing 0.25 μg/ml Hoechst No. 33342 (Sigma-Aldrich). Images were acquired using a DeltaVision CORE widefield fluorescence system fitted with a 60× NA 1.4 PlanApochromat objective (Olympus) and CoolSNAP charge-coupled device (CCD) camera (Roper Scientific). The microscope was controlled and images processed by SoftWorX acquisition and deconvolution software (GE Healthcare). All images are single, deconvolved optical sections.

### Fluorescence Two-Hybrid (F2H) Screens

For the F2H assays, a plasmid expressing mCherry-NLS-LacR-PP1ɣ was co-transfected with plasmids expressing GFP-tagged c20orf27 (wild type or KAGA mutant) in a HeLa cell line that contains an integrated lacO array[81]. Negative controls included co-expression of mCherry-NLS-LacR-PP1ɣ with free GFP, and co-expression of GFP-tagged regulatory proteins with mCherry-NLS-LacR. DNA was stained with Hoechst33342 and cells imaged live to monitor co-localization of mCherry and GFP signals.

### Bimolecular Fluorescence Complementation (BiFC) Assays

For Bimolecular Fluorescence Complementation (BiFC) assays[85], PP1γ fused to the N-terminal half of YFP (aa 1-154) was co-transfected in HEK293 cells with c20orf27 (wild type or KAGA mutant) fused to the C-terminal half of YFP (aa 155-238). DNA was stained with Hoechst33342 and the intensity and localization of EYFP fluorescence assessed by live imaging.

## Acknowledgments

We thank Dr. John Copeland and colleagues in the Trinkle-Mulcahy lab for helpful discussions and suggestions, and Himani Patel for contributions to the study. We also thank Drs. Richard Klemke, Anthony Hyman and Gregory Matera for reagents and Lawrence Puente at the Ottawa Hospital Research Institute Proteomics Core Facility for technical support.

## Author contributions

V.M. and L.T.-M. planned the experiments. V.M., D.C., J.L., S.O., D.C. and V.N. performed the experiments. V.M., G.B.M, F.-M.B. and L.T.-M. contributed intellectually to the work as well as read and edited the final manuscript.

## Data and materials availability

All data needed to evaluate the conclusions in the paper are present in the paper or the Supplementary Materials. Plasmids and cell lines are available through Addgene or upon request.

## Supplementary Materials

**Figure S1.**
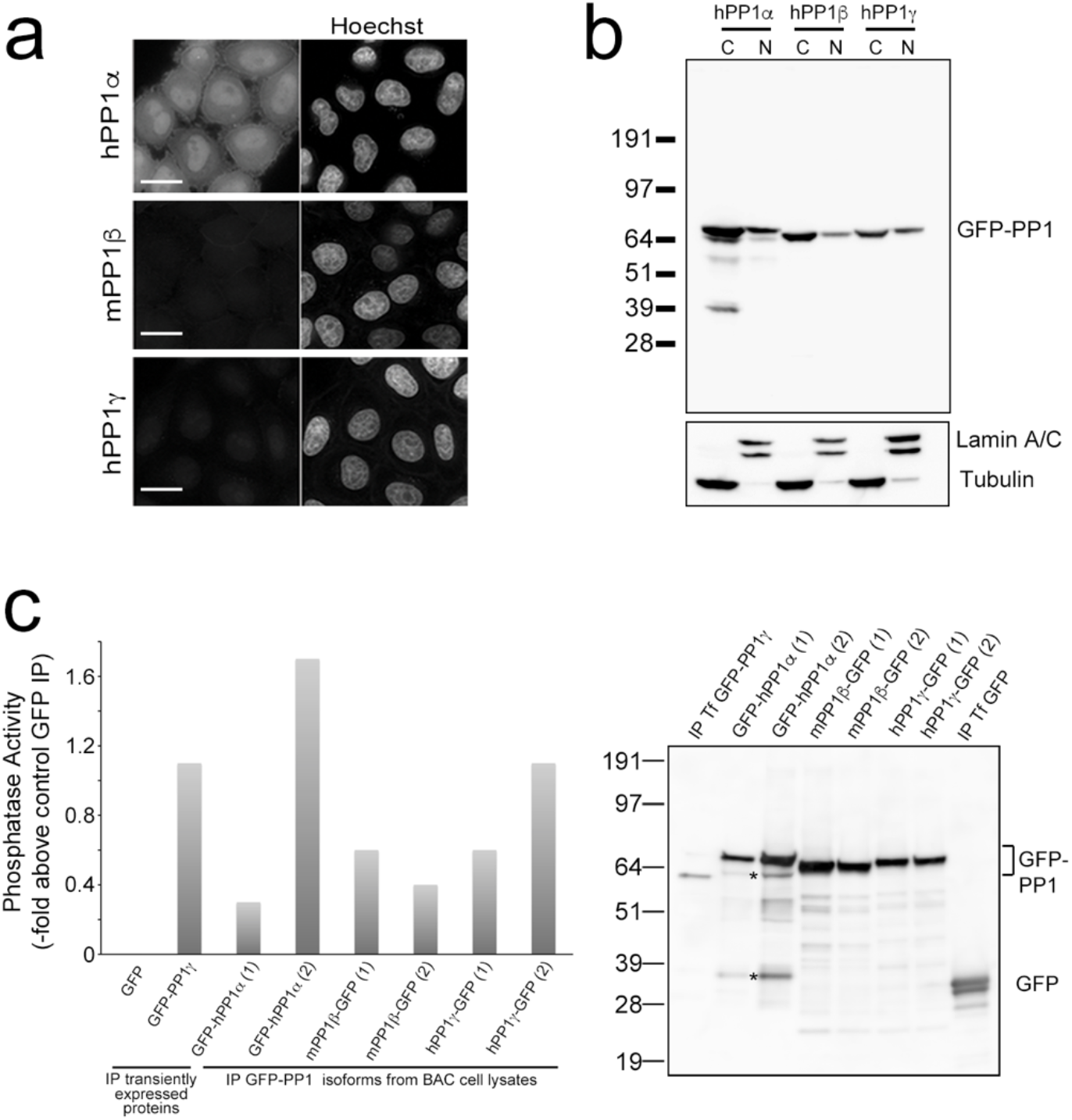
PP1ɑ is the most highly expressed isoform in HeLa/BAC stable cell lines. **(a)** Live imaging of GFP fluorescence (left panels) and Hoechst-stained DNA (right panels) in the 3 HeLa/BAC cell lines. Optimal image acquisition settings were determined for the GFP-PP1ɑ cell line and then the same exposure times (0.2 sec for GFP, 0.05 sec for Hoechst) were used to image the PP1β-GFP and GFP-PP1ɣ cell lines (bottom 2 rows). Scale bars are 5 µm. **(b)** Western blot analysis of GFP-PP1 isoforms with anti-GFP antibodies in cytoplasmic **(c)** and nuclear (N) fractions, using alpha-tubulin and Lamin A/C antibodies to detect these compartment-specific markers.

**Figure S2.**
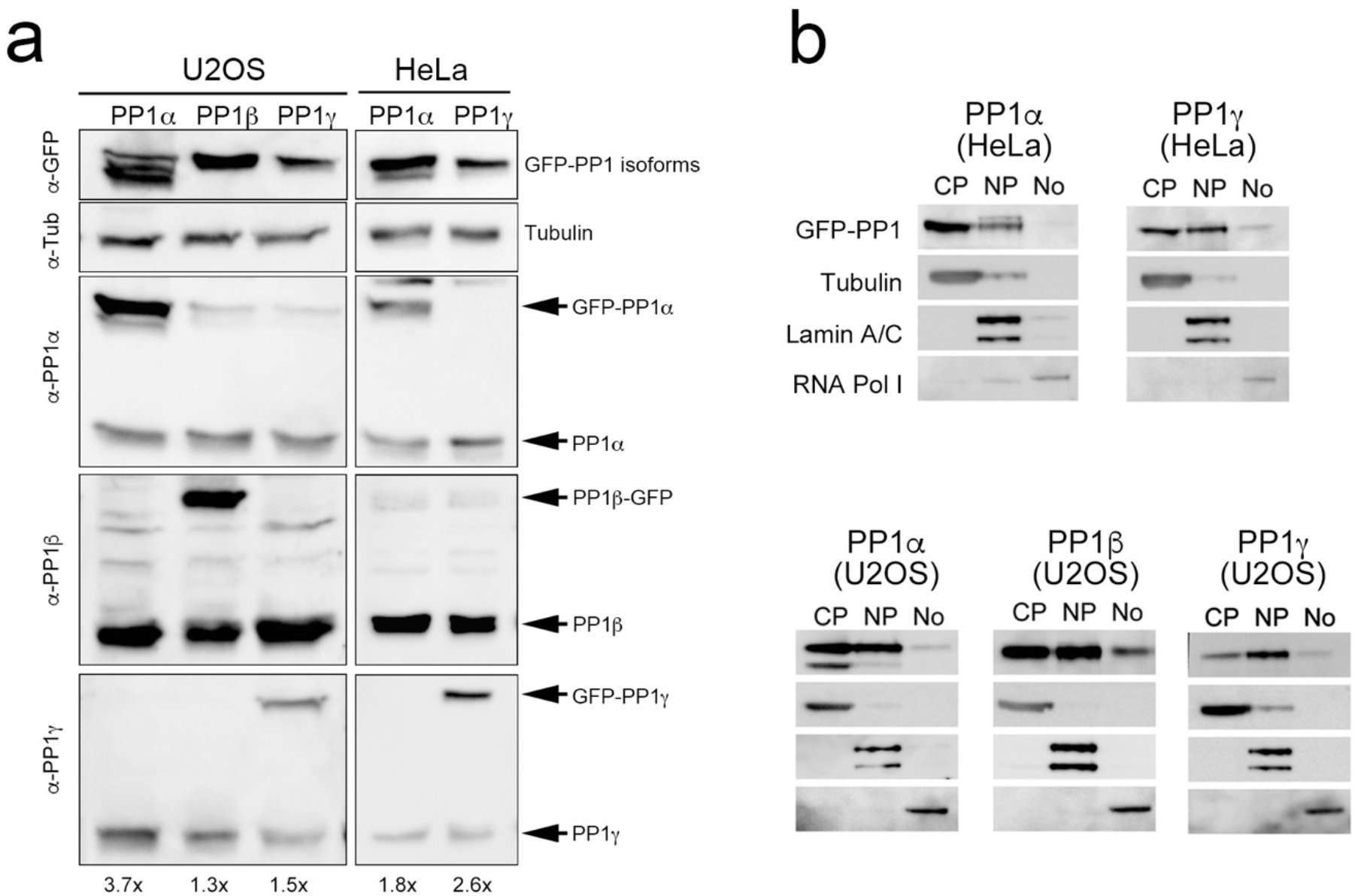
Expression of GFP-PP1 isoforms in HeLa and U2OS stable cell lines. **(a)** Whole cell extracts from the U2OS and HeLa stable cell lines (created by random stable integration) were probed with either anti-GFP (and anti-tubulin as a loading control) or with the isoform-specific PP1 antibodies. The approximate fold-level of expression of each fusion protein above the endogenous protein is indicated at the bottom. **(b)** Western blot analysis of cell equivalent volumes of cytoplasmic (CP), nucleoplasmic (NP) and nucleolar (No) extracts from the 5 stable cell lines, using markers for the 3 cellular compartments and anti-GFP to assess the distribution of the fusion protein between them.

**Figure S3.**
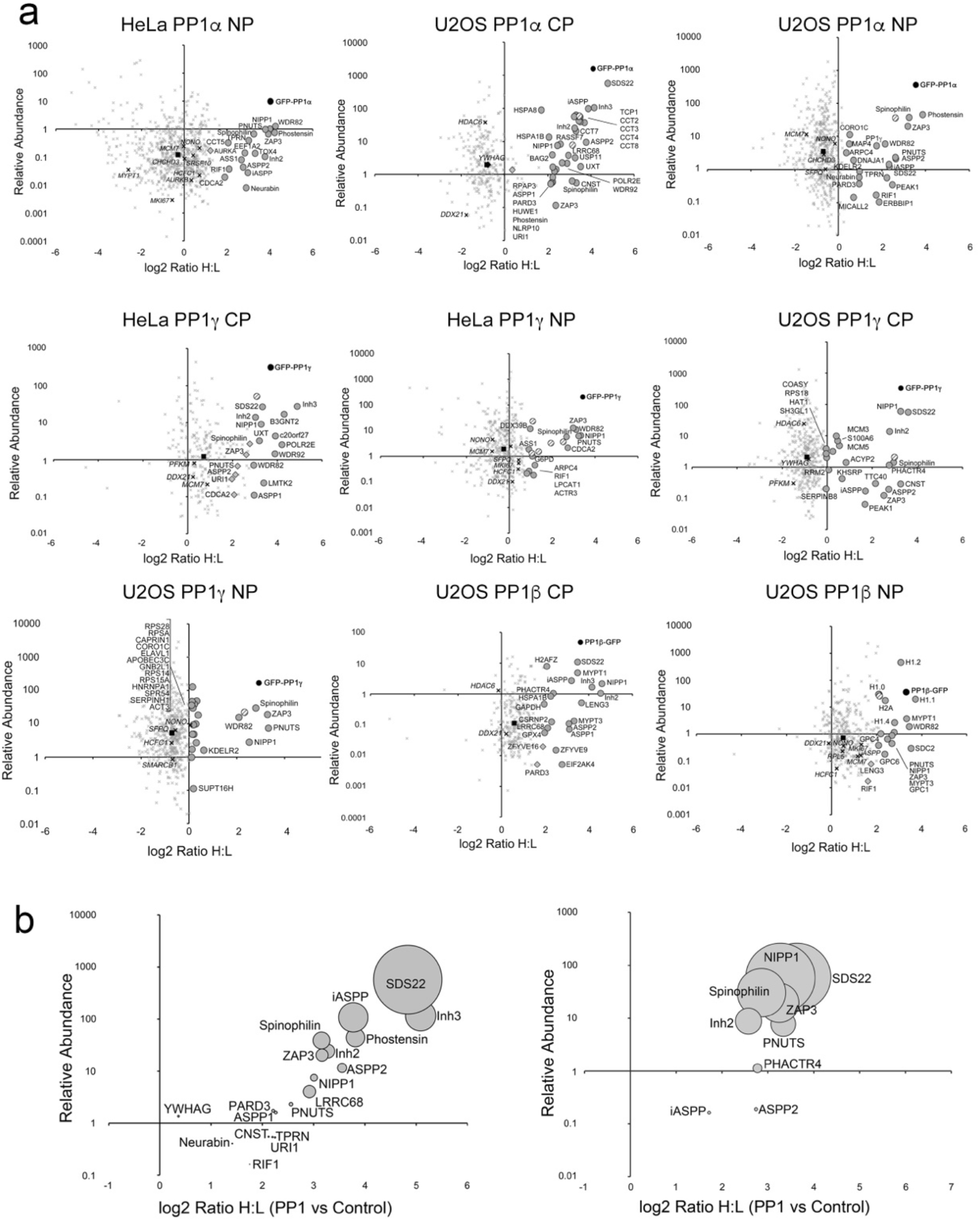
High confidence interactome datasets for GFP-PP1 isoforms stably expressed in HeLa and U2OS cells. **(a)** Cytoplasmic (CP) and Nucleoplasmic (NP) datasets were plotted as relative abundance vs. log2 Ratio H:L. Contaminant proteins captured non-specifically in both pulldowns cluster around a median value close to 0 (black square), while GFP-PP1 isoforms are highly enriched (black circle). Proteins with Significance B values < 0.05 are indicated (gray circles), as are known regulatory proteins that were enriched > 2-fold (gray diamonds). A black x marks known PP1 regulatory proteins (in italics) that were found at or below the cluster of contaminants, indicating that they likely bound non-specifically in this case. Hashed circles indicate proteins that were enriched in the control GFP dataset. **(b)** Bubble graphs visualizing the relative representation of known regulatory proteins in the U2OS GFP-PP1ɑ (left) and GFP-PP1ɣ (right) datasets.

**Figure S4.**
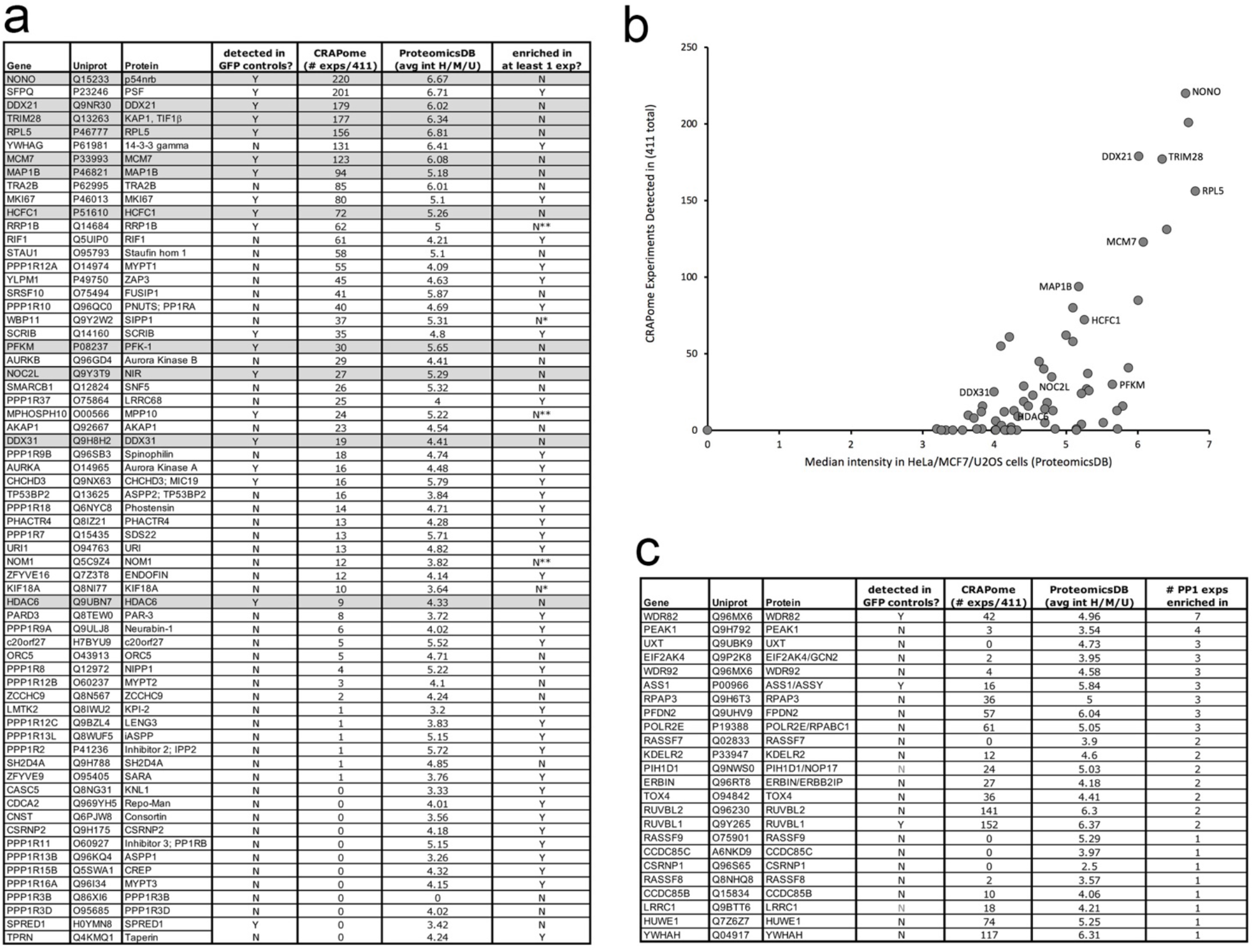
Some known regulatory proteins are common contaminants in AP/MS experiments. **(a)** Annotation of all known PP1 regulatory proteins identified in our screens (Table 1), noting whether or not they were detected in a GFP control experiment and/or enriched in at least one interactome screen. Further annotation includes the number of CRAPome contaminant experiments in which they were identified (out of a total of 411 deposited in the online repository; http://crapome.org) and their estimated expression level (log10 normalized iBAQ intensity) in HeLa, MCF7 and U2OS cells as annotated in the ProteomicsDB online repository for whole proteome datasets (http://proteomicsdb.org). An * indicates mitosis-specific interactors and ** indicates nucleolar interactors. **(b)** Plotting the CRAPome vs. ProteomicsDB information highlights abundant known regulatory proteins with a higher likelihood of binding non-specifically in AP/MS experiments. Those detected in our GFP controls and not enriched in at least one isoform dataset are noted on the graph (gray highlighting in A). **(c)** A similar analysis was carried out for the high confidence hits in Table 2, adding the number of individual interactome experiments in which they were enriched.

Table S1. List of primary and secondary antibodies and dyes used for Western blot analysis and immunofluorescence.

Supplemental Data File 1. U2OS CP/NUC fractionation dataset.

Supplemental Data File 2. HeLa/BAC and MCF7 PP1 isoform interactome datasets.

Supplemental Data File 3. PP1ɑ datasets.

Supplemental Data File 4. PP1β datasets.

Supplemental Data File 5. PP1ɣ datasets.

Supplemental Data File 6. Mitotic datasets.

Supplemental Data File 7. GFP control datasets.

Supplemental Data File 8. c20orf27 interactome dataset.

